# Structure Elucidation, Biosynthesis and Biological Evaluation of Neosorangicin A, a Member of the Sorangicin Family

**DOI:** 10.64898/2026.01.26.701680

**Authors:** Franziska Fries, Sebastian Walesch, Rolf Jansen, Kristin von Peinen, Luisa Mehr, Linda Pätzold, Sabrina Karwehl, Kathrin Mohr, Andreas M. Kany, Ronald Garcia, Jörg Haupenthal, Theresia Stradal, Markus Bischoff, Marc Stadler, Rolf Müller, Jennifer Herrmann

## Abstract

Antimicrobial resistance represents an escalating global health crisis, with drug-resistant infections predicted to cause up to 10 million deaths annually by 2050, underscoring the urgent need for novel antibiotics. Natural products play a crucial role in the discovery and development of antibiotics, with myxobacteria emerging as a particularly promising source due to their ability to produce structurally diverse and bioactive compounds. One prominent example of antibiotics from myxobacteria are the sorangicins, potent inhibitors of the bacterial RNA polymerase (RNAP). Here, we report the isolation of two unprecedented compounds, neosorangicin A (**1**) and neosorangioside A (**2**), from *Sorangium cellulosum* strain Soce439, elucidated their molecular structures, thereby revealing significant structural variation in comparison to sorangicin, and describe their biosynthetic pathway. Neosorangicin A (**1**) exhibited strong activity against various Gram-positive bacteria, with enhanced potency on intracellular *Staphylococcus aureus*. In a murine wound infection model, a head-to-head comparison of neosorangicin A (**1**) and sorangicin A (**3**) provided useful insights into how the altered physicochemical properties, arising from the shortened side chain and the lack of the free carboxylic acid of neosorangicin A, influence the *in vivo* efficacy of sorangicin derivatives.

## INTRODUCTION

Global health has improved significantly since the golden age of antibiotic discovery. However, the continuing misuse and overuse of antimicrobials in humans, animals and plants accelerated the development of drug-resistant pathogens.^1^ Although antimicrobial resistance (AMR) is a natural evolutionary process, the alarming pace at which drug-resistant pathogens are emerging poses a global threat to public health.^2^ The consequences of this crisis are devastating with almost 5 million deaths being associated with bacterial AMR in 2019, emphasizing the need for immediate action to combat this threat.^3^ In particular, innovative and effective antibiotics are required; however, most pharmaceutical companies abandoned antibiotic research and development (R&D) due to economic reasons years ago, resulting in a lack of newly approved antibiotic classes with novel modes of action, and an insufficient R&D pipeline.^4^ The World Health Organization (WHO) calls for attention to the insufficiency in novel approaches in the pipeline for new antibacterials to effectively tackle the emergence and spread of drug-resistant infections, especially those caused by bacteria outlined in their recently updated bacterial priority pathogens list.^5,6^

Myxobacteria with their capability to produce a wide range of secondary metabolites continue to provide biologically active compounds with high chemical diversity and unprecedented modes of action.^7^ Exceptionally important are the highly active bacterial topoisomerase-inhibiting cystobactamids from *Cystobacter* sp.,^8,9^ the *Corallococcus coralloides* derived peptide antibiotic corramycin^10,11^ as well as corallopyronin A, an α-pyrone antibiotic currently in preclinical development for the treatment of filarial worm infections due to its high efficacy against their *Wolbachia* symbionts.^12,13^

Sorangicin A (**3**), first reported in 1985, represents a macrolide-polyether antibiotic from the myxobacterium *Sorangium cellulosum* with potent activity, especially against Gram-positive bacteria. The natural product inhibits the DNA-dependent RNA polymerase (RNAP), a validated target in antibacterial therapy. Despite the lack of apparent chemical or structural similarity, sorangicin A (**3**) shares the same RNAP β-subunit pocket as the first-line antituberculosis drug rifampicin and, similarly to the latter, inhibits transcription initiation. Notably, sorangicin A (**3**) exerts a distinct mechanism of inhibition on certain rifampicin-resistant *Mycobacterium tuberculosis* RNAPs, as it prevents the template-strand DNA from accessing the catalytic active site. This intriguing dual mechanism is attributed to its conformational flexibility, and highlights the potential of such adaptability in natural products as a basis for overcoming antibiotic resistance.^14–16^

Herein we report the identification of two variations of the sorangicin scaffold, neosorangicin A (**1**) and its glucoside neosorangioside A (**2**), from the myxobacterial strain *S. cellulosum* Soce439. The molecular structures of the two new derivatives were elucidated using extensive HRESIMS and NMR analyses, revealing a shortened side chain lacking the terminal carboxylic acid. Furthermore, we describe the biosynthetic pathway of neosorangicin A (**1**) and compare it to that of sorangicin A (**3**), discussing possible explanations for the distinctive structural features. The newly identified derivative (**1**) showed enhanced antimicrobial activity *in vitro* and promising intracellular activity against *S. aureus*. Ultimately, we describe the results of a head-to-head comparison of neosorangicin A (**1**) to sorangicin A (**3**) in an *in vivo* setting using a murine wound infection model.

## RESULTS AND DISCUSSION

### Isolation and Structure Elucidation

In our ongoing biological screening for antibiotics, strain *S. cellulosum* Soce439 stood out due to a strong antibacterial activity of its culture extracts. This was correlated to a new compound, neosorangicin A (**1**), by analytical-scale RP-HPLC fractionation and biological testing. The compound showed a characteristic UV absorption at 302 nm as a presumptive variant of the antibiotic sorangicin A (**3**) (C_47_H_66_O_11_, λ_max_ 301 nm). HPLC-HRESIMS of this compound indicated the elemental composition C_44_H_62_O_10_ for **1** by molecular ion clusters [M+H]^+^ at *m/z* 751.4426 and [M+Na]^+^ at *m/z* 773.4253 in the positive mode as well as a cluster [M+HCO_2_H-H]^−^ at *m/z* 795.4335 in the negative mode.

The structure of **1** was elucidated from 1D and 2D NMR data in CD_3_OD (Figure 1, Table S1). Comparison of the new NMR data with those of sorangicin A (**3**) revealed that not only the triene part expected from the UV spectrum but also the complete lactone and the basis of the side chain was retained in **1** (Tables S2 and S3). In neosorangicin A (**1**) the structural part of C-1 to C-3 of sorangicin A (**3**) was absent. C-4 constitutes the new chain end as a methyl group while C-5 was assigned as a new secondary alcohol. The resulting new chiral center was characterized using the chiral bidentate NMR-solvents *R*,*R*- and *S*,*S*-bis-1,3-methylbenzylamine-2-methylpropane (BMBA-*p*-Me, **4**) (Figure S1),^17,18^ revealing the *R*-configuration of the secondary alcohol by comparison of the ^13^C-chemical shift differences in *R*,*R*- and *S*,*S*-BMBA for C-4 and C-6. The ^13^C NMR signal of the new methyl group C-4 was easily identified and Δ*δ* was unambiguously determined as positive (+0.1 ppm). Although the ^13^C NMR signal of C-6 could be one of three signals between 38 and 41 ppm the sign of the shift difference was negative for either of the three signal pairs since Δ*δ* was −0.01, −0.04 and −0.08 ppm (Figure S2). Furthermore, H-5 displayed a small vicinal coupling with H-6 (*J*_5,6_ = 4.4 Hz), indicating a *gauche* conformation of these protons. This assignment was supported by correlations in the ^1^H,^1^H ROESY spectrum between the two protons.

**Figure 1.**
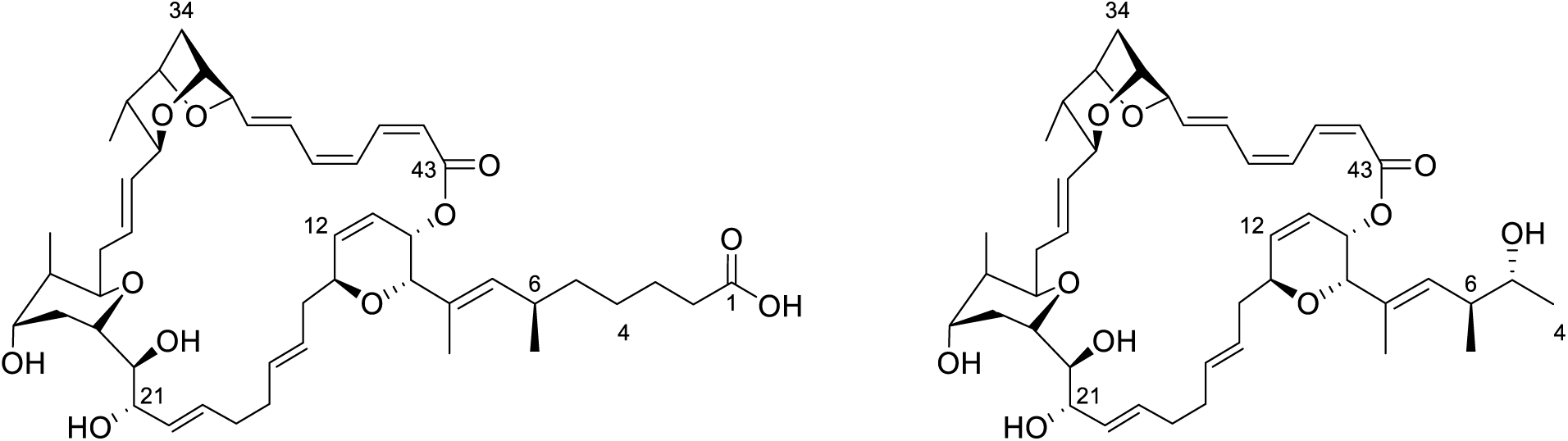
Chemical structures of sorangicin A (**3**, left) and neosorangicin A (**1**, right).

The production of neosorangicin A (**1**) further provided another, however weaker biologically active sorangicin derivative (**2**), which was initially recognized as such from its characteristic UV spectrum. In the HRESIMS the elemental composition C_50_H_72_O_15_ was derived from the molecular ion cluster [M+H]^+^ at *m/z* 913.4948 and [M-H]^−^ at *m/z* 911.4782. Additionally, a fragment with the elemental composition C_44_H_60_O_9_ was observed at *m/z* 733.4314 indicating the elimination of a hexose residue (C_6_H_12_O_6_). The NMR data showed that the aglycon of the glycoside **2** was neosorangicin A (**1**), which was glycosylated at position C-21, characterized by mutual ^1^H,^13^C HMBC correlations of methines C-21 and C-1’ (Table S4). The NMR data of the carbohydrate part were identical to those observed for sorangioside A (**4**), the β-_D_-glucopyranoside of sorangicin A (**3**) (Table S5). Consequently, **2** was identified as the β-_D_-glucopyranoside of neosorangicin A (**1**) and named neosorangioside A (Figure 2).

**Figure 2.**
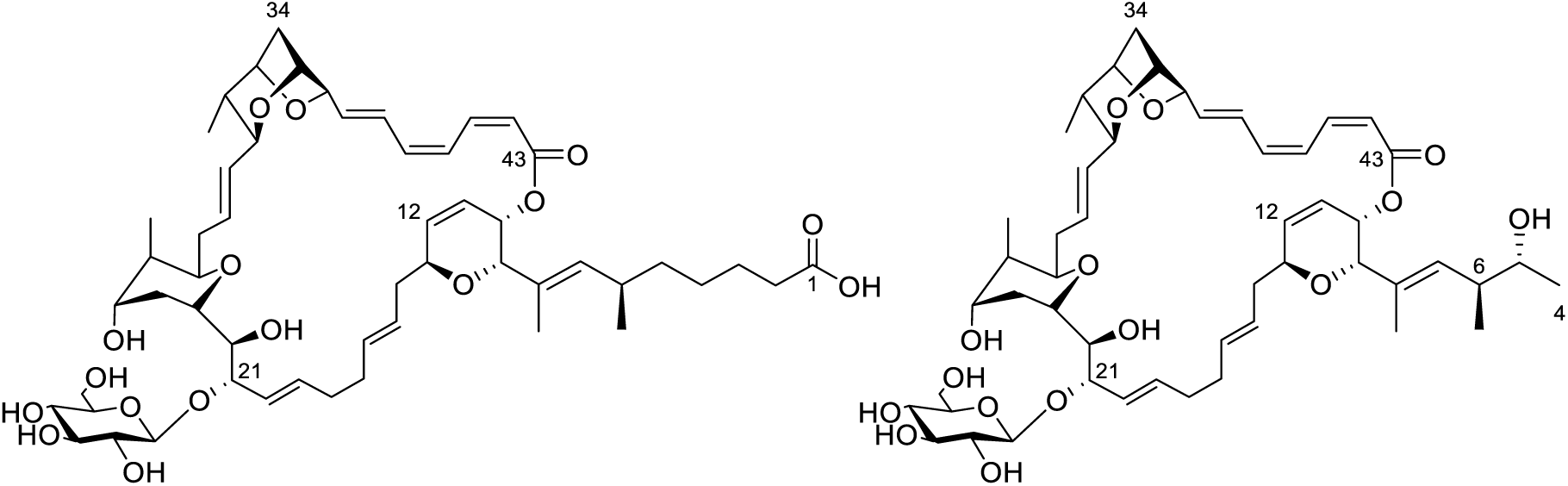
Chemical structures of sorangioside A (**4**, left) and neosorangioside A (**2**, right).

### Biosynthesis

As the genome sequence quality of the original producer *S. cellulosum* Soce439 was very poor, we interrogated our internal strain collection for further neosorangicin producers with sequenced genomes. Based on its high-quality genome sequence, we eventually chose the alternative producer Soce417 for the *in silico* analysis of the neosorangicin biosynthetic pathway.

The structural similarity of neosorangicin A (**1**) to sorangicin A (**3**) is also reflected by the high similarity of their biosynthetic gene clusters (Figure 3A). The neosorangicin biosynthetic gene cluster (*nsr* BGC) comprises 21 genes and spans 124.3 kb. Differently from the *sor* BGC in Soce12^19^, the biosynthetic machinery is encoded in seven instead of eight genes as *sorG* and *sorH* are condensed to *nsrGH*. Furthermore, the *nsr* BGC does not contain homologs of the putative amidase SorP^19^. All other core biosynthetic and surrounding genes in the *nsr* BGC display an identity of 83.2-94.7% on a protein level to their respective counterparts (Table S6). The high similarity of the *nsr* BGC to the *sor* BGC is further reflected in the architecture of the biosynthetic machinery as it features only a few differences between both pathways (Figure 3B, Table S7). Half of these differences are the occurrence of additional tandem acyl carrier protein (ACP) domains in modules 3, 5, 13 and 14 of the neosorangicin biosynthetic pathway. Such tandem ACP domains are believed to increase the efficiency of rate-limiting biosynthetic steps.^20^ Moreover, the biosynthetic machinery in the *nsr* BGC features an additional dehydratase (DH) domain in module 2, an additional ketoreductase (KR) domain in module 11 as well as an O-methyltransferase (O-MT) domain in module 18 and lacks a KR domain in module 1 compared to the sorangicin biosynthetic pathway.

**Figure 3.**
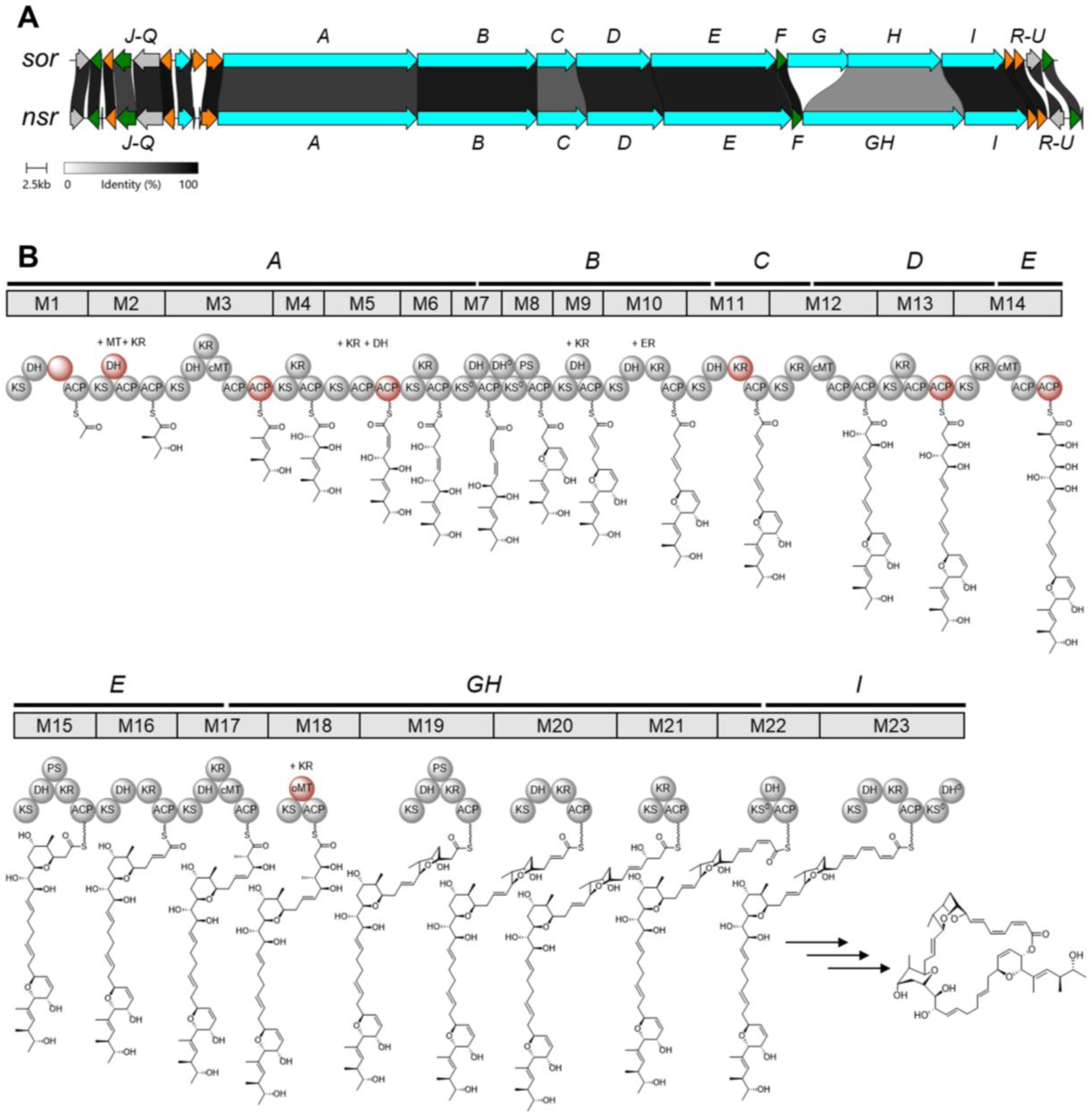
Distinct features of the *nsr* biosynthetic gene cluster (BGC). **A** Comparison between the *sor* BGC from Soce12 producing sorangicin A (**3**) and the *nsr* BGC from Soce417 producing neosorangicin A (**1**). Core biosynthetic genes are displayed in blue, additional biosynthetic genes in orange, resistance genes in green, other genes are coloured in grey. **B** Biosynthetic model for neosorangicin A, based on retro-biosynthesis, phylogenetic analysis of KS domains and occurrence of further domains within the respective modules. Domains that are missing from the sorangicin A BGC are coloured red, and the different architecture in module 1 of the *nsr* BGC is highlighted by an empty red circle. Putative *trans*-acting domains required to explain the observed structures are indicated above each module without bubbles. Superscripted “0” marks inactive domains. ACP, acyl carrier protein; DH, dehydratase; ER, enoylreductase; KR, ketoreductase; KS, ketosynthase; MT, methyltransferase; PS, pyran synthase. Phosphopantetheinyl arms are drawn as wavy lines.

Following the biosynthetic logic of multimodular megasynthases, the structural differences between **1** and **3** should be introduced within the first two modules of the neosorangicin assembly line. According to retro-biosynthetic considerations, module 1 of the *nsr* BGC should incorporate an acetyl intermediate, differing from the glutaryl intermediate in the sorangicin biosynthesis. However, bioinformatic analysis of the ketosynthase (KS) domains in module 1 of the *nsr* and *sor* BGCs with the transATor tool^21^ places them in the same clade of KS domains with a substrate specificity for unusual and larger starter units (Table S7). This classification was further supported by the high similarity of 94.0% of both KS domains in pairwise alignment (Figure S4). Consequently, similar to the initiation of the biosyntheses of **3** and sorangicin P^22^, the start of the biosynthesis of **1** remains speculative. Furthermore, the incorporation of an acetyl-intermediate in module 1 seems to contradict the predicted substrate specificity of the KS domain in module 2, as, similar to the respective KS in the *sor* BGC, it should mainly accept growing polyketides that are fully reduced at the β-position (Table S7, Figure S4). However, as the intermediate that was generated in module 1 does not possess a β-position yet, it might be accepted as a substrate by the KS domain of module 2. Another possibility may be the incorporation of a glutaryl-intermediate in module 1 of the neosorangicin biosynthesis with a loss of C-1 to C-3 in a later stage in or after the biosynthesis. The latter theory seems to be rather unlikely, as it would involve several enzymes that are encoded elsewhere in the genome and to our knowledge, such a post-assembly line modification has not been reported.

As the KS in module 3 accepts α-methylated substrates, that are either fully saturated or bear keto or _D_-hydroxy residues, it seems very likely that the growing ketide is methylated in module 2 by an *in trans*-acting C-methyltransferase (cMT). Following previous studies with trans-AT cMTs, this methylation should yield an *R*-configured intermediate^23^, corresponding to the observed stereochemistry of **1** and **3**. Considering the additional and possibly active DH domain in module 2 of the *nsr* BGC that would eliminate a putative hydroxyl-residue at C-5 to form an olefin, which in turn would contradict the predicted substrate specificity of KS 3 (Table S7) as well as retro-biosynthetic considerations, an *in trans* ketoreduction without a subsequent *in trans* enoyl reduction of the formed olefin in this module seems unlikely. Therefore, the hydroxyl-residue at C-5 should be either formed through an oxidation of the formed methylene or by the reduction of the unaffected ketone by a distinct enzyme at a later stage of the biosynthesis. As the *nsr* BGC does not encode any oxidoreductases or proteins with unknown functions, the origin of the hydroxyl-residue at C-5 remains elusive.

While in the biosynthesis of sorangicin A (**3**) an *in trans*-acting KR domain was hypothesized to be part of the reductive loop in module 11^19^, the respective module in the *nsr* BGC features a KR domain, agreeing with retro-biosynthesis and the predicted substrate specificity of KS 12 (Table S7). Provided an *in trans* ketoreduction of the ketide at C-33 in module 18 of the neosorangicin biosynthetic pathway, methylation of the resulting alcohol through the additional oMT domain in this module is in agreement with the predicted substrate specificities of the KS domains in module 19 of the *nsr* (Table S7), *sor* and *srb* BGCs.^19,22^ In the mature natural product C-33 is part of a 5-membered ring in the dioxabicyclooctane moiety, but similarly to the biosyntheses of sorangicin A (**3**) and sorangicin P, the formation of the ether between C-33 and C-36 cannot be fully explained through the biosynthetic machinery.^19,22^ If it is formed with the involvement of an elsewhere in the genome encoded, multifunctional cytochrome P450 monooxygenase that oxidizes C-36, as hypothesized in the sorangicin biosynthesis^19^, the original residue of C-33 could be replaced during ether formation. In analogy to sorangicin C as a minor product of the *sor* BGC, the *nsr* BGC should form derivatives that feature the C-31 to C-35 instead of the dioxabicyclooctane moiety. As neither homologs of sorangicin C, nor their methylated counterparts could be detected in the culture extracts of Soce417, it remains elusive if the O-MT domain in module 18 is active or not.

Similar to the previously described sorangicin BGCs, the *nsr* BGC does not contain a thioesterase (TE) domain to explain the chain release and macro-lactone formation of the neosorangicins. As discussed for the biosynthesis of sorangicin A, these reactions could be catalyzed by one of the amidohydrolases NsrK, NsrR or NsrS, by NsrM or NsrT, which are of unknown function or by a thioesterase that is encoded elsewhere in the genome.^19^

In accordance with the published ability of the glycosyltransferase SorF to form sorangioside A (**4**) from sorangicin A (**3**)^19,24^, it seems very likely that neosorangicin A (**1**) is glycosylated by NsrF to form neosorangioside A (**2**).

### Biological characterization

In light of sorangicin A’s (**3**) potent antibacterial activity, we sought to assess the bioactivity of neosorangicin A (**1**) using a panel of ESKAPE pathogens (Table 1). Compound **1** showed excellent inhibitory activities, in particular against Gram-positive pathogens with minimum inhibitory concentrations (MICs) in the mid ng mL^−1^ and low µg mL^−1^ range. Strikingly, **1** showed > 10-fold reduced MIC values against (methicillin-resistant) *S. aureus* compared to its congener **3**. Furthermore, activities against *Enterococci* were improved, and both derivatives show comparable potencies against the high-priority pathogen *M. tuberculosis*. In contrast, neosorangioside A (**2**) displayed only some weak activity on Gram-positive indicator strains (MICs of 16.6 µg mL^−1^ against *M. luteus* DSM1790 and *S. aureus* DSM346, respectively). Similar to **3** and rifampicin, **1** is less active on Gram-negative pathogens. Interestingly, the antimicrobial activity against *E. coli* could be significantly increased when sub-inhibitory concentrations of polymyxin B nonapeptide (PMBN) were added, which leads to a permeabilization of the outer membrane. In addition, the MIC values of **1** and **3** against efflux-deficient *E. coli* (TolC as part of the AcrAB multidrug efflux system) were decreased. This leads to the conclusion that the lower activity of sorangicins **1** and **3** against Gram-negative pathogens is mainly caused by insufficient uptake and partly also by efflux.

**Table 1.**
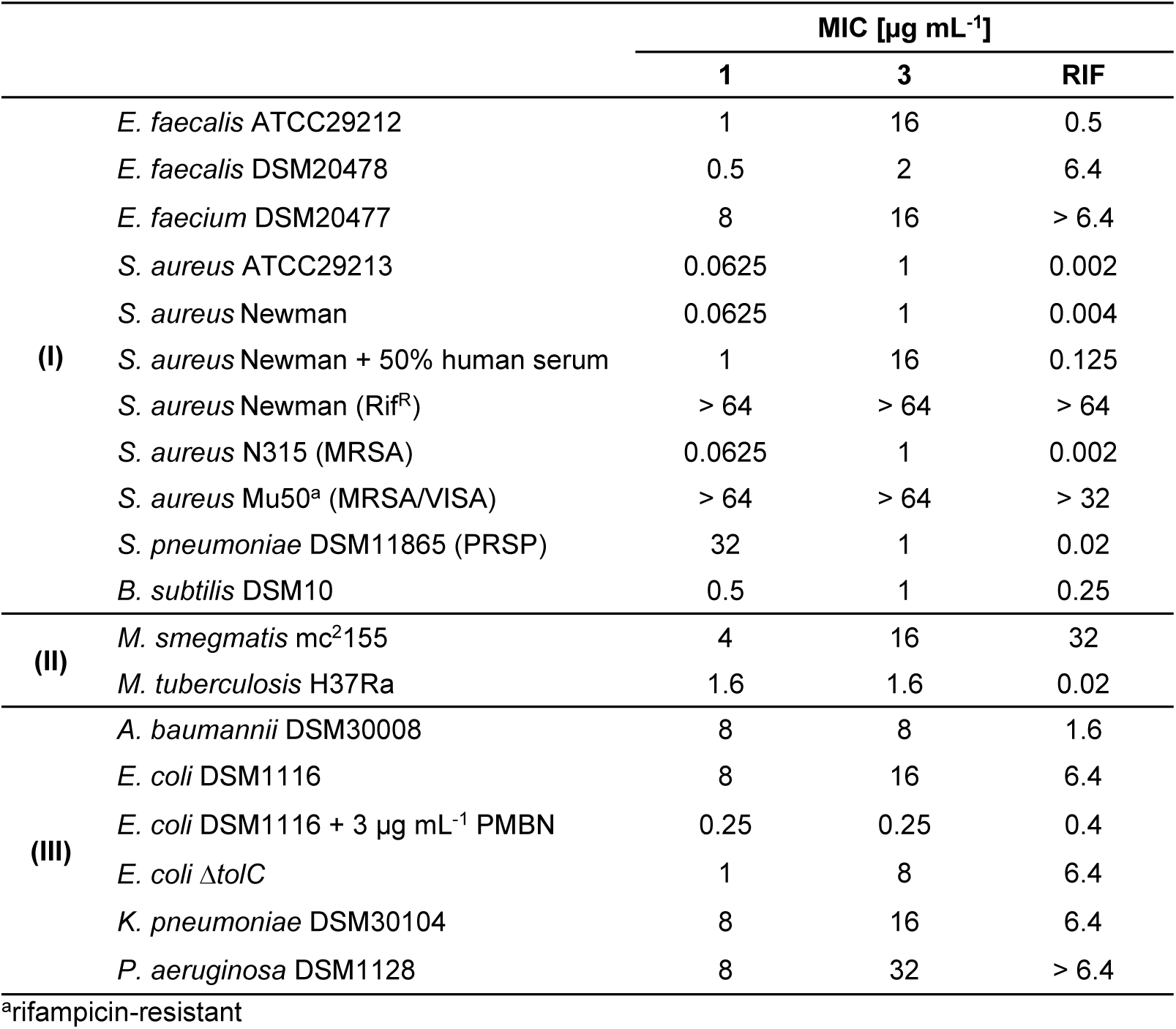
Minimum inhibitory concentrations (MICs) of neosorangicin A (**1**) and sorangicin A (**3**) on selected Gram-positive bacteria (I), mycobacteria (II) and Gram-negative (III) species. MRSA, methicillin-resistant *Staphylococcus* aureus; PMBN, polymyxin B nonapeptide; PRSP, penicillin-resistant *S. pneumoniae*; RIF, rifampicin; VISA, vancomycin-intermediate *S. aureus*.

Additionally, we determined IC_50_ values with purified *S. aureus* RNAP to confirm on-target activity of the newly identified sorangicin. The IC_50_ values were 0.08 µg mL^−1^, 0.31 µg mL^−1^, and 0.04 µg mL^−1^ for sorangicin A (**3**), neosorangicin A (**1**) and rifampicin, respectively (Figure S5). The increased activity of neosorangicin A (**1**), especially on *S. aureus*, might thus be contributed to a better cellular uptake rather than improved target binding on RNAP. Testing of **1** and **3** on rifampicin-resistant (Rif^R^) *S. aureus* Newman resulted in loss of activity (Table 1), confirming overlapping binding sites on RNA polymerase.

Recently, it has become accepted in the field that *S. aureus* can invade and replicate within host cells, representing a major reservoir for chronic and relapsing staphylococcal infections.^25,26^ This prompted us to determine the intracellular activity of neosorangicin A (**1**). Human-derived A549 cells and murine fibroblasts NIH 3T3 were infected with *S. aureus* Newman at a multiplicity of infection (MOI) of 100 and, following elimination of extracellular bacteria, exposed to different concentrations of neosorangicin A (**1**), sorangicin A (**3**), or rifampicin (Figure 4). To ensure that the antibiotics and solvents used have no adverse effects on murine or human cells, cytotoxicity and effects on proliferation were probed in their presence (Figure S6). Both sorangicin derivatives effectively reduced the intracellular *S. aureus* load at 1x MIC by more than one log_10_ unit, demonstrating efficacy comparable to rifampicin at the tested concentration (Figure 4). In human A549 cells, significant activity was also observed at sub-MIC concentrations (Figure 4A). Notably, **1** displayed approximately 10-fold higher intracellular potency than **3**, achieving the same reduction in bacterial load at 0.1 µg/mL as sorangicin A (**3**) at 1 µg/mL, consistent with its superior activity against extracellular bacteria (Figure 4 and Table 1).

**Figure 4.**
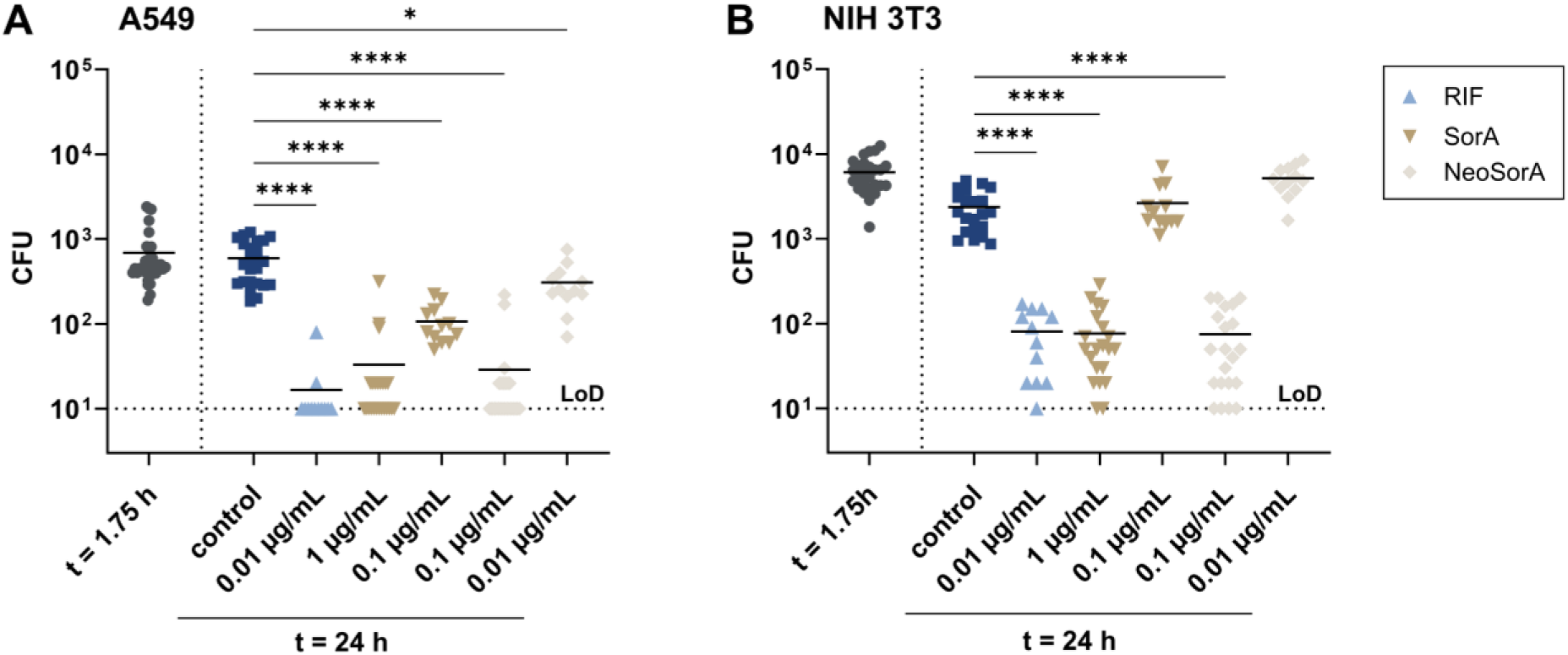
Intracellular activity of sorangicins against *S. aureus* Newman in A549 (**A**) and NIH 3T3 (**B**) cells. At 1.75 hours post infection, cells were washed once and treated with different concentrations of rifampicin (RIF), sorangicin A (SorA), neosorangicin A (NeoSorA), or with 0.1% (*v/v*) methanol (control) in the presence of 10 µg mL^−1^ gentamicin. Reduction of bacterial burden was evaluated 24 hours post infection by CFU (colony-forming unit) counting. Limit of detection (LoD) is 10^1^ CFU. Horizontal lines represent mean values of at least three independent biological experiments. Statistical analysis was performed by one-way ANOVA with Dunnett’s multiple comparison test. p < 0.05: *, p < 0.0001: ****.

Encouraged by the promising intracellular activity against *S. aureus*, we set out to evaluate the *in vivo* potency of neosorangicin A (**1**) in comparison to sorangicin A (**3**). To this end, we applied an *S. aureus*-based wound infection model in hairless Crl:SKH1-Hrhr (SKH1) mice that was previously developed and successfully used for the assessment of RNAP inhibitors.^27^ Skin wounds in mice were generated via punch biopsy and wounds were subsequently infected with *S. aureus* Newman. We then performed topical treatment with either neosorangicin A (**1**), sorangicin A (**3**) or rifampicin, and studied the effects of the antibiotics on wound healing as well as body weight kinetics. In addition, we determined the remaining CFUs in the wounds at the end of the experiment. As the body weight of the mice is influenced by the number of living bacteria in the wounds, it can be used as an indicator for disease progression, in particular in the first days after infection (Figure 5A). However, systemically available drug after topical administration (not quantified) can also have an influence on body weights. While the mice with infected and sham-treated wounds lost 10.8 ± 2% of their body weight within 48 hours after infection, the weight loss in the antibiotic treatment groups was clearly reduced. Sorangicin A (**3**) had the most prominent effect with a significantly reduced body weight loss (*p* < 0.05) to 6.3 ± 1.4%. Rifampicin also showed a strong, yet non-significant, effect on body weight loss that is comparable to that of sorangicin A (**3**). It is worth mentioning that the administered dose of rifampicin was considerably lower (20-fold) than the applied dose of sorangicin A (**3**), and that higher doses would very likely lead to even stronger reductions in body weight loss. The weight loss in the neosorangicin A (**1**) group amounted to 6.8 ± 4.4%, however the rather high standard deviation hindered a reliable evaluation of the drug. The wound healing provided a clearer picture for neosorangicin A (**1**) in which the drug was not capable of accelerating wound closure as the progression of wound closure was just as slow as for infected and sham-treated mice. This is accompanied by high bacterial counts of approximately 10^5^ CFU recovered from the wounds of mice at day 14 (Figures 5B and C). For sorangicin A (**3**), on the other hand, we observed significantly accelerated wound healing that was comparable to the uninfected wounds. Rifampicin also led to a significant change in the speed of wound healing, yet the change was not as pronounced as for sorangicin A (**3**). Both drugs, however, led to a significant reduction in bacterial burden of the infected wounds with >2-log fold reduction compared to sham-treated mice at day 14, underlining their strong antibacterial effect *in vivo* when administered topically.

**Figure 5.**
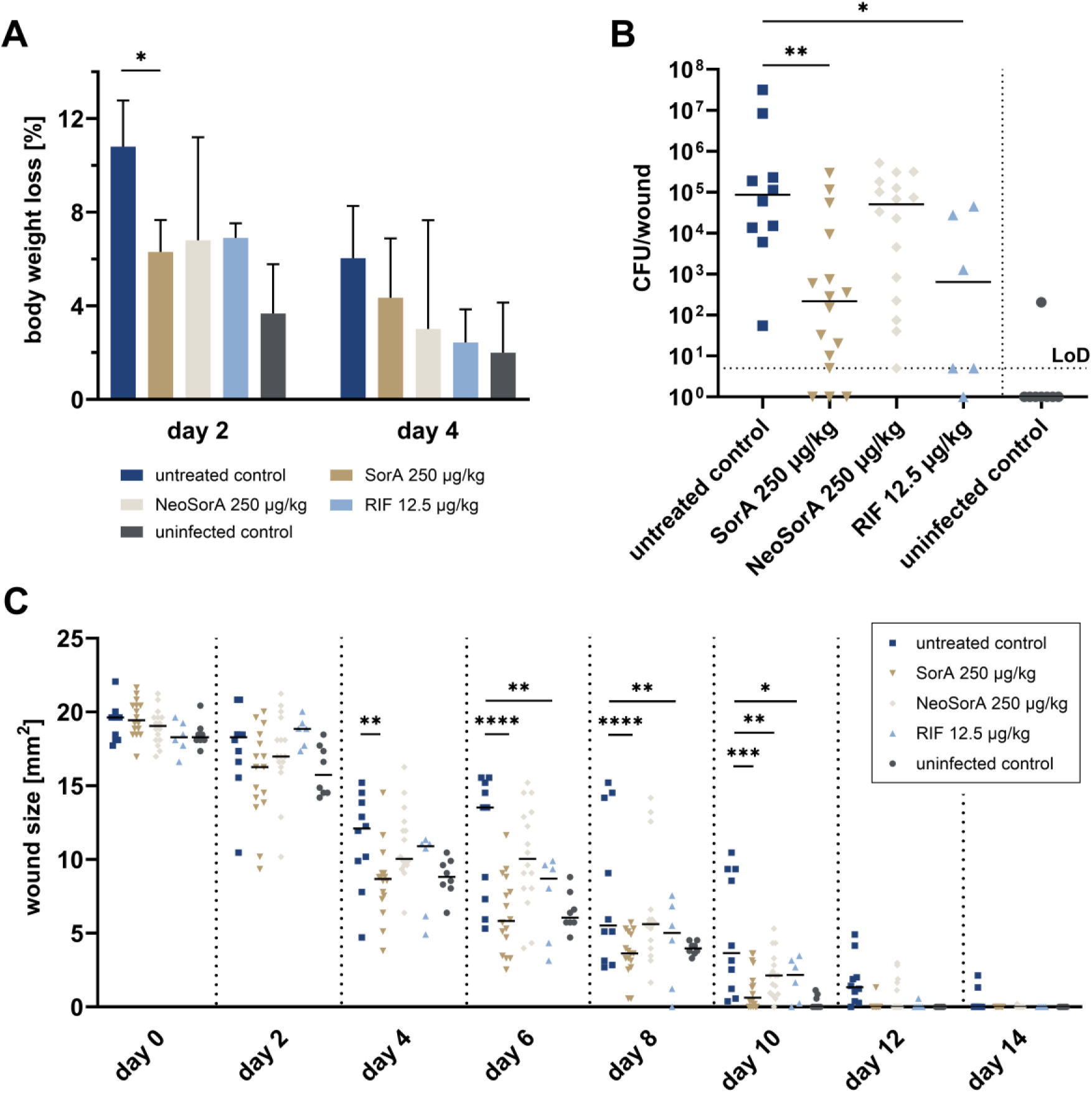
Evaluation of sorangicins in a *S. aureus-*based skin wound infection model in SKH1 mice. In this model, punch wounds of 5 mm in diameter were induced on both flanks of the mouse, and wounds were subsequently infected with 10^5^ CFU of *S. aureus* strain Newman. Infected wounds were treated either with sorangicin A (SorA; 250 µg/kg body weight), neosorangicin A (NeoSorA; 250 µg/kg body weight) or rifampicin (RIF; 12.5 µg/kg body weight) at 3 h, 48 h and 96 h post infection. **A** Loss of body weight (as % of starting weight) at 48 h and 96 h post infection. Mean values with standard deviations are illustrated. **B** Total numbers of CFU per wound in each treatment group (14 days after infection). Horizontal lines represent median values. Limit of detection (LoD) is 5×10^0^ CFU/wound. **C** Median wound areas (in mm^2^) over 14 days. Untreated control represents wounded and infected mice that were treated with the vehicle only, whereas uninfected control represents wounded but not infected mice. Asterisks indicate a significant difference. Statistical analysis was performed by two-way ANOVA with Dunnett’s multiple comparison test (for body weight loss and wound size) and by Kruskal-Wallis test with Dunn’s multiple comparisons test (for CFU numbers). *p* < 0.05: *; *p* < 0.01: **; *p* < 0.001: ***, *p* < 0.0001: ****. CFU: colony forming unit.

Despite its enhanced *in vitro* potency, neosorangicin A (**1**) was ineffective in eradicating *S. aureus in vivo*. Given the strong antibacterial efficacy of its relative sorangicin A (**3**) in the applied wound infection model, and the high structural similarity, with the only differences located in the side chain, we hypothesize that the structural variations, which significantly alter physicochemical properties and also result in changes of kinetic properties of neosorangicin A (**1**), render **1** ineffective *in vivo*. Since wound exudate is derived from plasma, the two fluids share similar compositions.^28,29^ Therefore, compounds with *e.g.*, low plasma stability or high plasma protein binding (PPB) are expected to display similar characteristics in wound exudate. *In vitro* ADME profiling revealed that neosorangicin A (**1**) is highly unstable in mouse plasma, with a plasma half-life of less than 0.5 min – considerably shorter than that of sorangicin A (**3**), which itself is known to be rapidly degraded. This pronounced instability in plasma, and likely in wound exudate, may account for its lack of *in vivo* activity. Notably, like **3**, **1** shows species-dependent variability in plasma clearance (Table 2). Thus, it cannot be excluded that neosorangicin A (**1**) may demonstrate *in vivo* efficacy in models employing other species.

**Table 2.** *In vitro* ADME of neosorangicin A (**1**). ADME data of sorangicin A (**3**) is adapted from ^30^. t_1/2_: half-life; CL_int_: intrinsic clearance; PPB: plasma protein binding. ^a^could not be determined due to rapid degradation in mouse plasma.

| Species | Metabolic stability |  | Plasma stability<br>t<sub>1/2</sub> [min] | PPB [%] |
| --- | --- | --- | --- | --- |
|  | Liver microsomes t<sub>1/2</sub> [min] | CL<sub>int</sub> [μL/mg/min] |  |  |
| Neosorangicin A |  |  |  |  |
| Mouse | 0.8 ± 0.3 | 1998 ± 755 | < 0.5 | —<sup>a</sup> |
| Rat | 0.6 ± 0.1 | 2405 ± 239 | 78.0 ± 6.0 | 96.9 ± 1.0 |
| Human | 1.6 ± 0.4 | 949 ± 261 | > 240 | 83.6 ± 5.0 |
| Sorangicin A |  |  |  |  |
| Mouse | 3.4 ± 2.6 | 557 ± 297 | 17.5 ± 7.1 | 87.9 ± 4.8 |
| Rat | 4.4. ± 0.5 | 239 ± 155 | > 240 | 88.8 ±1.7 |
| Human | 15.7 ± 2.8 | 90.4 ± 16.5 | > 240 | 87.0 ± 0.4 |

## CONCLUSION

Myxobacteria have emerged as a valuable source for structurally diverse and pharmacologically active metabolites, showcasing high potential to address the gaps in the drug discovery pipeline.^7,31^ In this study, we introduce two new members of the sorangicin family: neosorangicin A (**1**) and its glucoside (**2**), which were isolated through an activity-guided process from the myxobacterial strain *S. cellulosum* Soce439. Unlike the previously identified sorangicin P, which features structural modifications in the macrolactone ring,^22^ neosorangicin A (**1**) retains the same cyclic core structure as sorangicin A (**3**), differing only in its side chain, which is shorter and contains a secondary alcohol while lacking the terminal carboxylic acid. In accordance with the high structural similarities between neosorangicin A (**1**) and sorangicin A (**3**), the *nsr* BGC is very similar to the previously reported *sor* BGC. Although the structural differences between both sorangicin sub-classes are incorporated via PKS modules 1-3, they display a very similar architecture and show identical substrate specificity predictions. While the macrolactone ring in **1** is not different from the one in **3**, the *nsr* BGC features additional domains in modules 11 and 18 that correspond better to the observed structures or the predicted substrate specificities of the following KS domains in comparison to the *sor* BGC.

Despite the high structural similarity, neosorangicin A (**1**) demonstrates enhanced efficacy against several pathogenic Gram-positive bacteria with the activity extending to intracellular *Staphylococci*. Intrigued by the superior *in vitro* activity, we hoped for improved translatability of **1** and thus, as a next step, aimed to compare neosorangicin A (**1**) to sorangicin A (**3**) *in vivo*. However, while **3** showed outstanding *in vivo* activity, **1** failed to produce a significant effect in the applied model, likely due to rapid degradation in wound fluid. We thus conclude that neosorangicin A does not qualify as an improved starting point for antibiotic development. These results highlight the importance of determining ADME/PhysChem properties early in the drug discovery process and not solely rely on *in vitro* potency for the selection of promising derivatives as critical liabilities may be overlooked that can impede costly subsequent (pre)clinical translation.

## EXPERIMENTAL SECTION

### General experimental procedure

Optical rotations were determined with a Perkin-Elmer 241 instrument, UV data were recorded on a Shimadzu UV/Vis-2450 spectrophotometer in methanol (UVASOL, Merck. ^1^H NMR and ^13^C NMR spectra were recorded on Bruker Ascend 700 NMR spectrometers, locked to the deuterium signal of the solvent. Data acquisition, processing, and spectral analysis were performed with standard Bruker software and ACD/NMRSpectrus. Chemical shifts are given in parts per million (ppm), and coupling constants in hertz (Hz). HRESIMS data were recorded on a MaXis ESI TOF mass spectrometer (Bruker Daltonics), and molecular formulas were calculated including the isotopic pattern (Smart Formula algorithm). Analytical RP HPLC was carried out with an Agilent 1260 HPLC system equipped with a diode-array UV detector (DAD) and a Corona Ultra detector (Dionex) or a MaXis ESI TOF mass spectrometer (Bruker Daltonics). HPLC conditions: XBridge C_18_ column 100×2.1 mm (Waters), 3.5 μm, solvent A: H_2_O/acetonitrile (95/5), 5 mmol NH_4_Ac, 0.04 mL/L CH_3_COOH; solvent B: H_2_O/acetonitrile (5/95), 5 mmol NH_4_Ac, 0.04 mL/L CH_3_COOH; gradient system: 10% B increasing to 100% B in 30 min; flow rate 0.3 mL/min; 40 °C.

### Cultivation of *Sorangium cellulosum* strain Soce439

The strain was stored at −80 °C. It was reactivated in 20 mL of liquid medium consisting of 0.5% soy peptone, 0.2% yeast extract, 0.1% MgSO_4_×7H_2_O, 0.1% CaCl_2_×2H_2_O, 8 mg/L Na-Fe-EDTA and 10% Glucose×7H_2_O. The culture was scaled up to 1 L and used as inoculum for a fermentation of strain Soce439 that was performed in the same medium as above but supplemented with 2% Amberlite XAD-16 resin in a 70 L bioreactor. The bioreactor was kept at 30 °C, aerated at 0.05 vvm per minute and agitated with a flat blade turbine stirrer at 100 rpm for 384 h while the pH was regulated at 7.2-7.4. At the end of fermentation the XAD resin (1.71 kg) was recovered from the culture broth by sieving.

### Isolation and purification of neosorangicins

The XAD adsorber resin was extracted in a glass column with 2 L of methanol/H_2_O (3/7) and 3 L of methanol. The methanol extract was evaporated to an aqueous mixture, diluted with water, and extracted with ethyl acetate (three times with 350 mL). The combined ethyl acetate was dried with Na_2_SO_4_. It was evaporated to give 5.3 g of crude extract. The crude extract was resolved in 250 mL methanol with 1% H_2_O and extracted three times with heptane (three times 250 mL). The methanol was evaporated to give 3.91 g of an enriched crude extract. The crude extract was dissolved in methanol and filtered two times by a Strata-column (10 g, 55 µm, 70 Å). The methanol was evaporated to give 3.45 g of crude extract. The crude extract was separated by RP-MPLC [column 480×30 mm (Kronlab), ODS-AQ C_18_, 15 µm; solvent A: methanol/H_2_O (1/1); solvent B methanol; gradient system: 30% B holding at 30% B for 5 min, increasing to 50% B in 135 min, increasing to 100% B in 20 min and holding at 100% B for 60 min; flow rate: 30 mL/min; UV detection at 300 nm]. Two fractions with pure neosorangicin A (**1**) (247 mg) and neosorangioside A (**2**) (166 mg) were obtained.

Neosorangicin A (**1**): C_44_H_62_O_10_, M = 750.96; [α]^D^_20_ = + 137.8 (c = 0.4, MeOH); TLC: R*_f_* = 0.34 (DCM/MeOH 9/1); UV (MeOH) λ_max_ (log ε): 302 (4.366) nm; NMR see Table S1; HRESIMS: *m/z* 795.4335 [M+HCOOH-H]^−^ (calcd for C_45_H_63_O_12_^−^, 795.4325); *m/z* 751.4612 [M+H]^+^ (calcd for C_44_H_63_O_10_^+^, 751.4416), *m/z* 773.4498 [M+Na]^+^ (calcd for C_44_H_62_NaO_10_^+^, 773.42385); *m/z* 733.4508 [M-H_2_O+H]^+^ (calcd for C_44_H_61_O_9_^+^, 733.4310); *m/z* 715.4399 [M-2H_2_O+H]^+^ (calcd for C_44_H_59_O_8_^+^, 715.4204).

Neosorangioside A (**2**): C_50_H_72_O_15_, M = 913.10; [α]^D^_20_ = +108.4 (c = 0.5, MeOH); TLC: R*_f_* = 0.07 (DCM/MeOH 9/1); UV (MeOH) λ_max_ (log ε): 302 (4.375) nm; NMR see Table S2; HRESIMS: *m/z* 957.4837 [M+HCOOH-H]^−^ (calcd for 957.54853 [C_50_H_72_O_15_+HCOOH-H]^−^); *m/z* 911.4782 [M-H]^−^ (calcd for 911.4798 [C_50_H_72_O_15_-H]^−^); *m/z* 913.4948 [M+H]^+^ (913.4943 calcd for [C_50_H_72_O_15_+H]^+^); *m/z* 935.4767 [M+Na]^+^ (935.4763 calcd for [C_50_H_72_O_15_+Na]^+^); *m/z* 733.4314 [M-C_6_H_12_O_6_+H]^+^ (733.4310 calcd for [C_44_H_60_O_9_+H]^+^).

### Cultivation of *S. cellulosum* Soce417, extraction and data analysis for analytical purposes

*S. cellulosum* Soce417 was grown in triplicate in 300 mL shake flasks containing 50 mL CyH medium (0.3% casitone, 0.15% yeast extract, 0.8% soluble starch, 0.2% soy flour, 0.2% D-glucose, 0.1% MgSO_4_×7H_2_O, 0.1% CaCl_2_×2H_2_O and 8 mg/L Na-Fe-EDTA) inoculated with 5% (*v/v*) pre-culture. The medium was supplemented with 4% (*v/v*) of a sterile aqueous solution of XAD-16 adsorber resin (Sigma Aldrich) to bind secondary metabolites in the culture medium. After 10 days of cultivation, the cultures were pelleted in 50 mL falcon tubes in an Eppendorf centrifuge at 5804R at 8,288 *g* and 4 °C for 10 min. The pellets were then freeze-dried and subsequently extracted with 40 mL MeOH and stirred at 250 rpm at room temperature for 3 h. The supernatants were decanted into round flasks through 125-µm folded filters. The solvent and potential residual water were removed on a rotary evaporator with a water bath temperature of 40 °C, at appropriate pressures. The dried extracts were dissolved/resuspended in 1000 µL MeOH and stored at −20 °C until further analysis. For the purpose of UHPLC-hrMS analysis, the crude extracts were diluted 1:3 with methanol and centrifuged at 21,500 *g* and 4 °C (HIMAC CT15RE, Koki Holdings Co.) for 5 min to remove residual insolubilities such as salts, cell debris, and XAD fragments.

UPLC-MS measurement were performed on a Dionex (Germering, Germany) Vanquish Flex UHPLC system equipped with Waters (Eschborn, Germany) BEH C18 column (100 × 2.1 mm, 1.7 μm) equipped with a Waters VanGuard BEH C18 1.7 μm guard column. Separation of 1 µL sample was achieved by a linear gradient from (A) H_2_O + 0.1% FA to (B) ACN + 0.1% FA at a flow rate of 600 µL/min and 45 °C. The gradient was initiated by a 0.5 min isocratic step at 5% B, followed by an increase to 95% B in 18 min to end with a 2 min step at 95% B before re-equilibration with initial conditions. UV-vis spectra were recorded by a DAD in the range from 200 to 600 nm. The timsTOF fleX was operated in positive ESI mode, with 1.0 bar nebulizer pressure, 5.0 L/min dry gas, 200 °C dry heater, 4000 V capillary voltage, 500 V end plate offset, 500 Vpp funnel 1 RF, 250 Vpp funnel 2 RF, 80 V deflection delta, 5 eV ion energy, 10 eV collision energy, 1100 Vpp collision RF, 5 µs pre pulse storage, 65 µs transfer time. TIMS delta values were set to −20 V (delta 1), −120 V (delta 2), 80 V (delta 3), 100 V (delta 4), 0 V (delta 5), and 100 V (delta 6). The 1/K0 (inverse reduced ion mobility) range was set from 0.55 Vs/cm^2^ to 1.87 Vs/cm^2^, the mass range was *m/z* 100-2000. MS^2^ spectra were acquired using the PASEF DDA mode with a collision energy gradient based on ion mobility: Starting at 25 eV for 0.55 Vs/cm^2^ (1/K0) to 35 eV at 1.2 Vs/cm^2^ to 40 eV at 1.5 Vs/cm^2^ to 60 eV for 1.87 Vs/cm^2^. Ion charge control (ICC) was enabled and set to 7.5 Mio. counts. The analysis accumulation and ramp time was set at 100 ms with a spectra rate of 9.43 Hz and a total cycle of 0.32 sec was also selected resulting in one full TIMS-MS scan and two PASEF MS/MS scans. Precursor ions were actively excluded for 0.1 min and were reconsidered if the intensity was 2.0-fold higher than the previous selection with a target intensity of 4000 and an intensity threshold of 100. TIMS dimension was calibrated linearly using 4 selected ions from ESI Low Concentration Tuning Mix (Agilent Technologies, USA) [*m/z*, 1/k0: (301.998139, 0.6678 Vs/cm^2^), (601.979077, 0.8782 Vs/cm^2^)] in negative mode and [*m/z*, 1/k0: (322.048121, 0.7363 Vs/cm^2^), (622.028960, 0.9915 Vs/cm^2^), (922.0098, 0.9915 Vs/cm^2^), (622.028960, 0.9915 Vs/cm^2^)] i n positive mode. The mobility for mobility calibration was taken from the CCS compendium.^32^ Calibration was done automatically before every LC-MS run by injection of a basic sodium formate solution through a filled 20 µL loop switched into the LC flow at the beginning of each run.

### Genome sequencing of *S. cellulosum* Soce417 and characterization of biosynthetic pathway

Whole genome sequencing of the alternative neosorangicin producer *S. cellulosum* Soce417 was performed by Plasmidsaurus using their hybrid sequencing platform of Oxford Nanopore Technology and Illumina sequencing technology with custom analysis and annotation. This analysis yielded eight scaffolds with a total length of 16,026,845 bp that were analyzed with antiSMASH 8.0.1.^33^ The published sorangicin BGC from *S. cellulosum* Soce12 (GenBank accession code HM584908) was used for biosynthetic pathway comparison. Protein alignments were done in Geneious, version 2025.2.2 by the Geneious Alignment tool (default values).

### Antimicrobial susceptibility testing

Sorangicin derivatives were tested in microbroth dilution assays on a panel of Gram-positive and Gram-negative bacteria. All microorganisms were obtained from the German Collection of Microorganisms and Cell Cultures (Deutsche Sammlung für Mikroorganismen und Zellkulturen, DSMZ) and the American Type Culture Collection (ATCC) or were part of our internal strain collection. Overnight cultures were prepared from cryopreserved cultures and were diluted to achieve a final inoculum of 10^5^ CFU mL^−1^. Serial dilutions of compounds in DMSO were prepared in sterile 96-well plates in the test medium (cation-adjusted Müller-Hinton broth; supplemented with 2.5% (*v/v*) lysed horse blood for *Enterococcus* and *Streptococcus* spp.) The cell suspension was added and microorganisms were grown for 18-24 h at either 30 °C or 37 °C. Streptococci and enterococci were grown under microaerophilic conditions. Growth inhibition was assessed by visual inspection and given MIC values are the lowest concentration of antibiotic at which there was no visible growth. The same method was used for testing *Mycobacterium smegmatis*, but with the use of Middlebrook 7H9 complete medium supplemented with oleic acid, albumin, dextrose and catalase (OADC, 10%). *M. smegmatis* plates were incubated for 48 h at 37 °C. For assessing activity against *Mycobacterium tuberculosis*, an adapted resazurin microtitre assay (REMA) was performed as previously described.^34^ In short, *M. tuberculosis* single cells were prepared and added to compound dilutions in M7H9. Plates were incubated for 6 d at 37 °C, followed by addition of 50 µL of resazurin and incubation for another day at 37 °C. The MIC was determined visually and additionally confirmed by measuring fluorescence (excitation at 530 nm, emission at 590 nm).

### RNA polymerase inhibition

The inhibition of *S. aureus* RNA polymerase by Sorangicin derivatives **1** and **3** was tested *in vitro* using a bacterial RNA Polymerase Assay Kit (ProFoldin, MA, USA). The samples were prepared and analyzed according to the manufacturer’s protocol with slight modifications. In brief, serial dilutions (concentration range: 0.1 ng mL^−1^ – 10 µg mL^−1^) of rifampicin, sorangicin A (**3**) and neosorangicin (**1**) in black 384-well plates (low-volume, non-binding surface; Corning) were incubated for 1 h at room temperature with 25 nM RNA polymerase from *S. aureus* in assay buffer (42.5 mM HEPES, 42.5 mM NH_4_Cl, 2 mM DTT, 4 mM MgCl_2_, 0.005 % (*v/v*) Tween-20; pH 7.5) containing 0.5 mM NTPs and 1x DNA template. The reaction was monitored by incubation with a fluorescent probe for 5 min and measurement of the fluorescence intensity at 535 nm after excitation at 485 nm (Tecan Infinite M200PRO). Half-inhibitory concentrations (IC_50_) were determined by sigmoidal curve fitting.

### MTT cell cytotoxicity assay

Human alveolar epithelial A549 cells (ATCC CCL-185) and murine fibroblast NIH 3T3 cells (ATCC CRL-1658), obtained from the American Type Culture Collection, were cultured in DMEM with 4.5 g/L glucose, 10% fetal bovine serum, 2 mM _L_-glutamine, 1mM sodium pyruvate and 1% non-essential amino acids at 37 °C and 7.5% CO_2_. Cytotoxicity of the test compounds was assessed to determine half-maximal inhibitory concentrations (IC_50_). Cells were seeded at a density of 5×10^4^ cells/mL in 96-well plates and allowed to attach for 30 min at 37 °C and 7.5% CO₂. In parallel, compound dilutions were prepared in a separate 96-well plate, including Epothilone B as a positive control and methanol as solvent control. Cells were incubated with 60 µL of the respective compound dilutions for 5 days at 37 °C and 7.5% CO₂. After incubation, cell viability was assessed by adding MTT solution (5 mg mL^−1^) and incubating for 2 h. Following incubation, the formation of reduced MTT (formazan) was quantified using a microplate reader at 595 nm.

### Proliferation assay

To assess the effects of compounds on cell growth, 3×10^3^ A549 or NIH 3T3 cells were seeded onto a 96-well plate and incubated overnight at 37 °C and 7.5% CO₂. The next day, the compounds (0.1% (*v/v*) methanol, 1 µg mL^−1^ sorangicin A, 0.1 µg mL^−1^ neosorangicin A and 10 µg mL^−1^ of rifampicin) were added to the medium. Cells were imaged in quadruplicates using an Incucyte S3 live-cell analysis system (Sartorius, Göttingen, Germany) with a 20x objective at 37 °C and 5% CO_2_. Images of the phase contrast were captured at hourly intervals for a 24 hour period. The *Adherent Cell-by-Cell Analysis module* of the IncuCyte S3 Live Cell Analysis Software was used to quantify the object counts, and these were then normalized to the average of the initial phase contrast count.

### Intracellular efficiency

For infection experiments, A549 and NIH 3T3 cells were seeded in 24-well plates at a density of 5×10⁴ cells per well and incubated overnight to allow adherence. An overnight culture of *S. aureus* Newman (1:50) was inoculated into LB medium and grown at 37 °C under agitation until mid-log phase (OD_600_ 0.5–0.8). Bacteria were harvested by centrifugation at 5,000 rpm for 2 min and diluted in DMEM (without supplements) to obtain a MOI of 100. To infect the cells, the bacterial suspension was added directly to each well without removing the existing medium. Then, the plates were centrifuged at 2,000 rpm for 5 min to synchronize the infection. The infection proceeded for 90 min at 37 °C and 5% CO₂ followed by a 15 min incubation with DMEM containing 50 µg mL^−1^ gentamicin to eliminate extracellular bacteria. The cells were then washed once with PBS and incubated for up to 24 h post-infection (h.p.i.) in DMEM supplemented with 10 µg mL^−1^ gentamicin and the respective test compounds or methanol as a control (0.1% *v/v*): Rifampicin (0.01 µg mL^−1^), sorangicin A (1 µg mL^−1^ and 0.1 µg mL^−1^), and neosorangicin A (0.1 µg mL^−1^ and 0.01 µg mL^−1^). The cells were then washed three times with PBS, and the intracellular bacteria were recovered by lysing the cells with 0.5% Triton X-100 in PBS for five minutes at room temperature. The lysates were kept on ice to prevent further bacterial replication, serially diluted in PBS, and plated on LB agar. The intracellular bacterial load immediately after extracellular killing was determined by recovering intracellular bacteria in the same way before adding test compounds to serve as the post-gentamicin control (t = 1.75 h control). The plates were incubated overnight at 37 °C, and the colony-forming units (CFUs) were counted the following day.

### *In vitro* ADME studies

For the evaluation of phase I metabolic stability, neosorangicin A (1 μM) was incubated with 0.5 mg mL^−1^ pooled mouse (C57BL/6), Wistar rat or human liver microsomes (Xenotech, Kansas City, USA), 2 mM NADPH, 10 mM MgCl_2_ in 100 mM potassium phosphate buffer pH 7.4 at 37 °C for 120 min on a microplate shaker (Eppendorf, Hamburg, Germany). The metabolic stability of testosterone, verapamil and ketoconazole was determined in parallel to confirm the enzymatic activity of mouse/rat liver microsomes. For human liver microsomes, testosterone, diclofenac and propranolol were used. The incubation was stopped after defined time points by precipitation of aliquots of enzymes with 2 volumes of cold internal standard solution (15 nM diphenhydramine in 10% methanol/acetonitrile). Samples were stored on ice until the end of the incubation and precipitated protein was removed by centrifugation (15 min, 4 °C, 4,000 *g*). Remaining neosorangicin A at the different time points was analyzed by HPLC-MS/MS (Vanquish Flex coupled to a TSQ Altis Plus, Thermo Fisher, Dreieich, Germany) and used to determine half-life (t_1/2_).

To determine stability in plasma, neosorangicin A (1 µM) was incubated with pooled CD-1 mouse, Wistar rat or human plasma (Neo Biotech, Nanterre, France). Samples were taken at defined time points by mixing aliquots with 4 volumes of ice-cold internal standard solution (12.5 nM diphenhydramine in 10% methanol/acetonitrile). Samples were stored on ice until the end of the incubation and precipitated protein was removed by centrifugation (15 min, 4 °C, 4,000 *g*, 2 centrifugation steps). Remaining neosorangicin A at the different time points was analyzed by HPLC-MS/MS (Vanquish Flex coupled to a TSQ Altis Plus, Thermo Fisher, Dreieich, Germany). The plasma stability of procain, propantheline and diltiazem were determined in parallel to confirm the enzymatic activity.

Plasma protein binding was determined using the Rapid Equilibrium Dialysis (RED) system (Thermo Fisher Scientific, Waltham MA, USA). Neosorangicin A was diluted to 10 µM in 50% Wistar rat or human plasma (Neo Biotech, Nanterre, France) in PBS pH 7.4 and added to the respective chamber according to the manufacturer’s protocol, followed by addition of PBS pH 7.4 to the opposite chamber. Samples were taken immediately after addition to the plate as well as after 2, 4 and 5 h for human plasma and 50 min, 2 h and 4 h for rat plasma by mixing 10 µL with 80 µL ice-cold internal standard solution (12.5 nM diphenhydramine in 10% methanol/acetonitrile), followed by addition of 10 µL plasma to samples taken from PBS and vice versa. Samples were stored on ice until the end of the incubation and precipitated protein was removed by centrifugation (15 min, 4 °C, 4,000 *g*, 2 centrifugation steps). The amount of the remaining neorsorangicin A at the different time points was analyzed by HPLC-MS/MS (Vanquish Flex coupled to a TSQ Altis Plus, Thermo Fisher, Dreieich, Germany). The amount of neosorangicin A bound to protein was calculated using the equation PPB [%] = 100 – 100 x (amount in buffer chamber/amount in plasma chamber). For rat plasma, PPB is given at 50 min of incubation due to the observed degradation affecting overall recovery.

### *In vivo* mouse model

Animal experiments were performed with the approval of the local State Review Board of Saarland, Germany (project identification code 43/2016) and were conducted following the national and European guidelines for the ethical and human treatment of animals. All authors complied with the ARRIVE guidelines. Female SKH1 hairless mice (Crl:SKH1-Hr^hr^) were obtained from Charles River (Sulzfeld, Germany) and kept under specific pathogen-free conditions according to the regulations of German veterinary law. PBS-washed bacterial cells obtained from exponential growth phase cultures were used as inocula.

The *S. aureus*-based wound infection model was carried out essentially as described earlier.^27^ Briefly, seven to nine weeks old female hairless mice were anesthetized by intraperitoneal injection of 100 mg kg^−1^ body weight (bw) ketamine hydrochloride (Zoetis, Berlin, Germany) and 10 mg kg^−1^ bw of xylazine hydrochloride (Bayer, Leverkusen, Germany) and treated with a dose of carprofen (5 mg kg^−1^ bw, Zoetis, Berlin, Germany). After disinfection of the dorsal areas with ethanol (70%), full-thickness excisional punch wounds (Ø 5 mm) were created on both flanks through the skin down to the panniculus carnosus. Wounds were stabilized using silicone rings (HUG Technik und Sicherheit GmbH, Ergolding, Germany) and subsequently infected with 10 µL of a PBS suspension containing ∼10^7^ CFU mL^−1^ of *S. aureus* strain Newman. Infected wounds were allowed to dry for 5 min and were afterwards covered with Tegaderm (3M, Neuss, Germany). 10 µL aliquots of vehicle (10% DMSO, 10% Cremophor EL, 80% saline), sorangicin A (0.5 mg mL^−1^), neosorangicin A (0.5 mg mL^−1^), or rifampicin (25 µg mL^−1^) were spotted onto the infected wounds at 3 h, 48 h and 96 hours post infection. Body weights and wound sizes (measured with an electronic caliper [ChiliTec 17909, ChiliTec GmbH, Lehre-Essenrode, Germany] in mm with two decimal digits) were determined on every second day. After 14 days post infection, mice were euthanized by injecting a lethal dose of ketamine hydrochloride (250 mg kg^−1^ bw) and xylazine hydrochloride (25 mg kg^−1^ bw). After the absence of the inter-phalangeal reflex, the animal’s chest was opened and blood was taken by cardiac puncture. Afterwards, full-thickness tissues were harvested for microbial analyses. Excised wounds were homogenized in 0.5 mL PBS with a hand disperser (POLYTRON® PT 1200 E, Kinematica, Eschbach, Germany), and serial dilutions of the homogenates were plated on sheep blood agar (SBA) plates. CFU rates were determined after 24 h of cultivation at 37 °C. A blood stripe incubated on SBA at 37 °C served as an indicator of systemic bacterial infection.

## DATA AVAILABILITY

The identified neosorangicin (*nsr*) biosynthetic gene cluster was deposited in GenBank under the accession number XXXXXXXX (will be added upon acceptance of this manuscript). The raw NMR data files of neosorangicin A and neosorangioside A were deposited to nmrXiv and are accessible under XXXXXX (will be added upon acceptance of this manuscript).

## ACKNOWLEDGMENTS

We thank W. Collisi and K. Schober for excellent technical assistance. Furthermore, we thank C. Kakoschke for recording the NMR spectra, A. Gollasch for recording the HRESIMS spectra, and S. Bernecker and co-workers for large-scale cultivation. We acknowledge Nestor Zaburannyi for initial biosynthesis analysis.

