## Supplementary Information for "Structure Elucidation, Biosynthesis and Biological Evaluation of Neosorangicin A, a Member of the Sorangicin Family"

### Table of Contents


**Supplementary Tables** **3**

Table S1 3
Table S2 5
Table S3 6
Table S4 7
Table S5 9
Table S6 10
Table S7 11

**Supplementary Figures** **13**

Figure S1 13
Figure S2 13
Figure S3 13
Figure S4 14
Figure S5 15
Figure S6 16
Figure S7 17
Figure S8 18
Figure S9 19
Figure S10 20
Figure S11 21
Figure S12 22
Figure S13 23
Figure S14 24
Figure S15 25
Figure S16 26
Figure S17 27
Figure S18 28

**References** **29**

### Supplementary Tables

**Table S1.** NMR spectroscopic data of neosorangicin A (**1**) acquired at 700/175 MHz in methanol-$d_4$.

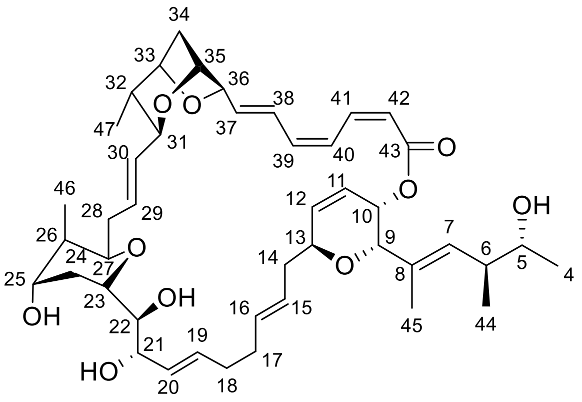

| Position | $\delta_C$[a] [ppm], type | $\delta_H$[b] [ppm], mult | COSY[c] | ROESY[d] | HMBC[e] |
|---|---|---|---|---|---|
| 4 | 19.3, CH3 | 1.01, d (6.5) | 5 | 5, 6, 7, 44 | 5, 6 |
| 5 | 71.8, CH | 3.56, qd (6.3, 6.3, 6.3, 4.5) | 4, 6 | 4, 6, 7, 44 | 4, 6, 7, 44 |
| 6 | 40.3, CH | 2.53, dqd (9.9, 6.9, 6.9, 6.9, 4.5) | 5, 7, 44 | 4, 5, 7, 44, 45 | 4, 5, 7, 8, 44 |
| 7 | 130.0, CH | 5.39, dquin (9.9, 1.3, 1.3, 1.3, 1.3) | 6, 9, 45 | 4, 5, 6, 9, 44 | 5, 6, 8, 9, 10, 44, 45 |
| 8 | 133.4, C | | | | |
| 9 | 74.3, CH | 4.25, br s | 7, 10, 45 | 7, 10, 14a, 45 | 7, 8, 10, 13, 45 |
| 10 | 66.9, CH | 5.32, dd (5.8, 1.7) | 9, 11 | 9, 11, 45 | 9, 11, 12, 43 |
| 11 | 123.9, CH | 6.03, ddd (9.9, 5.8, 2.2) | 10, 12, 13 | 10, 12 | 9, 10, 12, 13 |
| 12 | 137.1, CH | 6.14, dd (10.0, 3.1) | 11, 13 | 11, 13, 14b | 10, 11, 13 |
| 13 | 75.3, CH | 4.41, m | 11, 12, 14a, 14b | 12, 14b, 15 | 9, 11, 12, 14, 15 |
| 14a | 35.5, CH2 | 2.39, ddd (14.0, 10.7, 6.8) | 13, 14b, 15 | 9, 14b, 16 | 12, 13, 15, 16 |
| 14b | | 2.15, m | 13, 14a, 15 | 12, 13, 14a, 16 | 13, 15 |
| 15 | 128.5, CH | 5.53, m | 14a, 14b | 13, 17a, 17b | 14, 16 |
| 16 | 134.0, CH | 5.53, m | 17a, 17b | 14a, 14b, 18a, 18b | 15, 17, 18 |
| 17a | 33.9, CH2 | 2.15, m | 16, 18b | 15, 19 | 15, 16, 18, 19 |
| 17b | | 2.08, m | 16, 18a | 15, 18a, 19 | 15, 16, 18, 19 |
| 18a | 34.7, CH2 | 2.16, m | 17b, 19 | 16, 17b, 20 | 16, 17, 19, 20 |
| 18b | | 2.08, m | 17a, 19 | 16, 20 | 16, 17, 19, 20 |
| 19 | 134.6, CH | 5.72, ddd (15.0, 8.0, 5.6) | 18a, 18b, 20 | 17a, 17b, 21 | 18, 21 |
| 20 | 129.9, CH | 5.58, dd (15.4, 7.9) | 19, 21 | 18a, 18b, 21, 23 | 18, 19, 21, 22 |
| 21 | 74.5, CH | 4.16, dd (7.5, 4.3) | 20, 22 | 19, 20, 22 | 19, 20, 22, 23 |
| 22 | 77.6, CH | 3.47, dd (7.7, 4.3) | 21, 23 | 21, 24b | 20, 21, 23, 24 |
| 23 | 75.1, CH | 3.64, ddd (11.6, 7.7, 2.4) | 22, 24a, 24b | 20, 24a, 27 | 21, 22, 25, 27 |
| 24a | 31.1, CH2 | 1.73, m | 23, 24b, 25 | 23, 24b, 25 | 25, 26 |
| 24b | | 1.64, ddd (14.6, 11.6, 3.0) | 23, 24a, 25 | 24, 22, 24a, 25, 46 | 22, 23, 26 |
| 25 | 71.2, CH | 3.85, m | 24a, 24b, 26 | 24a, 24b, 26, 46 | 23, 26, 27, 46 |
| 26 | 38.3, CH | 1.55, m | 25, 27, 46 | 25, 27, 46 | 24, 25, 46 |
| 27 | 74.8, CH | 3.83, m | 26, 28a, 28b | 23, 26, 28a, 28b, 29 | 23, 25, 28, 29, 46 |
| 28a | 37.2, CH2 | 2.25, dddd (13.9, 5.6, 4.3, 1.7) | 27, 28b, 29 | 27, 28b, 29, 46 | 26, 27, 29, 30 |
| 28b | | 2.16, m | 27, 28a, 29 | 27, 28a, 30, 46 | 26, 27, 29, 30 |
| 29 | 133.0, CH | 5.45, ddd (15.0, 10.1, 4.1) | 28a, 28b, 30 | 27, 28a, 31 | 27, 28, 30, 31 |
| 30 | 133.0, CH | 5.37, ddd (15.2, 8.5, 1.8) | 29, 31 | 28b, 31, 32 | 28, 29, 31, 32 |
| 31 | 81.3, CH | 3.85, m | 30, 32 | 29, 30, 37, 38, 47 | 29, 30, 32, 33, 47 |
| 32 | 42.2, CH | 1.43, dq (9.4, 6.8, 6.8, 6.8) | 31, 33, 47 | 30, 33, 34b, 47 | 30, 31, 33, 34, 47 |
| 33 | 81.3, CH | 4.30, d (6.5) | 32, 34a | 32, 34a, 34b, 47 | 31, 34, 35, 36, 47 |

| | | | | | |
|---|---|---|---|---|---|
| 34a | 39.9, CH2 | 2.05, ddd (11.6, 6.6, 2.7) | 33, 34b, 35 | 33, 34b, 35, 36 | 32, 33 |
| 34b | | 1.92, dd (11.6, 1.3) | 34a, 35 | 32, 33, 34a, 35 | 32, 33, 35, 36 |
| 35 | 77.7, CH | 4.40, m | 34a, 34b, 36 | 34a, 34b, 36, 37 | 31, 33, 34, 36 |
| 36 | 82.2, CH | 4.59, br m | 35, 37, 38 | 34a, 35, 37, 38 | 37, 38, 39 |
| 37 | 135.6, CH | 6.24, dd (15.3, 3.9) | 36, 38, 39 | 31, 35, 36, 39 | 35, 36, 38, 39 |
| 38 | 127.5, CH | 7.03, ddd (15.2, 11.5, 1.5) | 36, 37, 39 | 31, 36, 41 | 36, 37, 39, 40 |
| 39 | 138.1, CH | 6.46, br dd (10.8, 9.7) | 37, 38, 40, 42 | 37, 40 | 37, 38, 41, 43 |
| 40 | 126.6, CH | 7.17, m | 39 | 39 | 38, 42, 43 |
| 41 | 139.2, CH | 7.16, m | 42 | 38, 42 | 38, 39, 42, 43 |
| 42 | 119.7, CH | 5.62, br d (9.7) | 39, 41 | 41 | 39, 40, 43 |
| 43 | 167.8, C | | | | |
| 44 | 15.7, CH3 | 0.91, d (6.9) | 6 | 4, 5, 6, 7 | 5, 6, 7 |
| 45 | 14.4, CH3 | 1.67, d (0.9) | 7, 9 | 6, 9, 10 | 6, 7, 8, 9, 10, 44 |
| 46 | 10.9, CH3 | 0.87, d (7.1) | 26 | 24, 25, 26, 28a, 28b | 25, 26, 27 |
| 47 | 15.6, CH3 | 0.83, d (6.9) | 32 | 31, 32, 33 | 31, 32, 33 |

[a] Acquired in methanol-$d_4$ at 176.1 MHz and calibrated to solvent signal at 49.2 ppm.
[b] Acquired in methanol-$d_d$ at 700.4 MHz and calibrated to solvent signal at 3.31 ppm.
[c] Proton showing COSY correlations to indicated proton.
[d] Proton showing ROESY correlations to indicated proton.
[e] Proton showing HMBC correlations to indicated carbon.

**Table S2.** $^{13}$C NMR comparison neosorangicin A (**1**) and sorangicin A (**3**) in methanol-$d_4$.

| Position | δ $^{13}$C [ppm] NeoA (1) | Position | δ $^{13}$C [ppm] SorA (3) | Δ (NeoA-SorA) |
|---|---|---|---|---|
| 4 | 19.3 | 4 | 28.2 | -8.9 |
| 5 | 71.8 | 5 | 38.5 | 33.3 |
| 6 | 40.3 | 6 | 33.0 | 7.3 |
| 7 | 130.0 | 7 | 134.2 | -4.1 |
| 8 | 133.4 | 8 | 131.2 | 2.2 |
| 9 | 74.3 | 9 | 74.4 | -0.1 |
| 10 | 66.9 | 10 | 66.9 | 0.0 |
| 11 | 123.9 | 11 | 123.8 | 0.1 |
| 12 | 137.1 | 12 | 136.9 | 0.2 |
| 13 | 75.3 | 13 | 75.3 | 0.0 |
| 14 | 35.5 | 14 | 35.5 | 0.1 |
| 15 | 128.5 | 15 | 128.3 | 0.1 |
| 16 | 134.0 | 16 | 133.6 | 0.4 |
| 17 | 33.9 | 17 | 33.4 | 0.6 |
| 18 | 34.7 | 18 | 34.0 | 0.8 |
| 19 | 134.6 | 19 | 134.4 | 0.3 |
| 20 | 129.9 | 20 | 130.2 | -0.3 |
| 21 | 74.5 | 21 | 74.4 | 0.1 |
| 22 | 77.6 | 22 | 77.8 | -0.1 |
| 23 | 75.1 | 23 | 75.1 | 0.0 |
| 24 | 31.1 | 24 | 30.9 | 0.3 |
| 25 | 71.2 | 25 | 71.1 | 0.1 |
| 26 | 38.3 | 26 | 38.5 | -0.2 |
| 27 | 74.8 | 27 | 74.9 | 0.0 |
| 28 | 37.2 | 28 | 37.1 | 0.1 |
| 29 | 133.0 | 29 | 133.0 | 0.0 |
| 30 | 133.0 | 30 | 132.8 | 0.2 |
| 31 | 81.3 | 31 | 81.2 | 0.1 |
| 32 | 42.2 | 32 | 42.2 | 0.1 |
| 33 | 81.3 | 33 | 81.0 | 0.2 |
| 34 | 39.9 | 34 | 39.9 | 0.0 |
| 35 | 77.7 | 35 | 77.6 | 0.1 |
| 36 | 82.2 | 36 | 82.3 | 0.0 |
| 37 | 135.6 | 37 | 134.9 | 0.7 |
| 38 | 127.5 | 38 | 127.8 | -0.3 |
| 39 | 138.1 | 39 | 137.6 | 0.5 |
| 40 | 126.6 | 40 | 127.0 | -0.4 |
| 41 | 139.2 | 41 | 139.1 | 0.1 |
| 42 | 119.7 | 42 | 119.7 | 0.0 |
| 43 | 167.8 | 43 | 167.7 | 0.2 |
| 44 | 15.7 | 44 | 21.7 | -6.0 |
| 45 | 14.4 | 45 | 14.3 | 0.1 |
| 46 | 10.9 | 46 | 10.9 | 0.0 |
| 47 | 15.6 | 47 | 15.4 | 0.3 |

**Table S3.** $^{1}$H NMR comparison neosorangicin A (**1**) and sorangicin A (**3**) in methanol-$d_4$.

| Position | δ $^{1}$H [ppm] NeoA (1) | Position | δ $^{1}$H [ppm] SorA (2) | Δ (NeoA-SorA) |
|---|---|---|---|---|
| 4a | | 4a | 1.36 | |
| 4b | 1.01 | 4b | 1.3 | -0.29 |
| 5a | 3.56 | 5a | 1.42 | 2.14 |
| 5b | 3.56 | 5b | 1.25 | 2.31 |
| 6 | 2.53 | 6 | 2.43 | 0.10 |
| 7 | 5.39 | 7 | 5.34 | 0.04 |
| 8 | | 8 | | 0.00 |
| 9 | 4.25 | 9 | 4.28 | -0.03 |
| 10 | 5.32 | 10 | 5.35 | -0.03 |
| 11 | 6.03 | 11 | 6.05 | -0.02 |
| 12 | 6.14 | 12 | 6.17 | -0.03 |
| 13 | 4.41 | 13 | 4.43 | -0.02 |
| 14a | 2.39 | 14a | 2.43 | -0.05 |
| 14b | 2.15 | 14b | 2.17 | -0.02 |
| 15 | 5.53 | 15 | 5.58 | -0.05 |
| 16 | 5.53 | 16 | 5.58 | -0.05 |
| 17a | 2.15 | 17a | 2.24 | -0.09 |
| 17b | 2.08 | 17b | 2.14 | -0.06 |
| 18a | 2.16 | 18a | 2.24 | -0.08 |
| 18b | 2.08 | 18b | 2.17 | -0.09 |
| 19 | 5.72 | 19 | 5.79 | -0.07 |
| 20 | 5.58 | 20 | 5.64 | -0.06 |
| 21 | 4.16 | 21 | 4.19 | -0.03 |
| 22 | 3.47 | 22 | 3.52 | -0.06 |
| 23 | 3.64 | 23 | 3.73 | -0.09 |
| 24a | 1.73 | 24a | 1.76 | -0.03 |
| 24b | 1.64 | 24b | 1.7 | -0.06 |
| 25 | 3.85 | 25 | 3.87 | -0.02 |
| 26 | 1.55 | 26 | 1.59 | -0.04 |
| 27 | 3.83 | 27 | 3.89 | -0.06 |
| 28a | 2.25 | 28a | 2.32 | -0.07 |
| 28b | 2.16 | 28b | 2.17 | -0.02 |
| 29 | 5.45 | 29 | 5.54 | -0.09 |
| 30 | 5.37 | 30 | 5.42 | -0.05 |
| 31 | 3.85 | 31 | 3.87 | -0.02 |
| 32 | 1.43 | 32 | 1.46 | -0.03 |
| 33 | 4.30 | 33 | 4.32 | -0.02 |
| 34a | 2.05 | 34a | 2.09 | -0.04 |
| 34b | 1.92 | 34b | 1.97 | -0.05 |
| 35 | 4.40 | 35 | 4.45 | -0.05 |
| 36 | 4.59 | 36 | 4.61 | -0.02 |
| 37 | 6.24 | 37 | 6.26 | -0.02 |
| 38 | 7.03 | 38 | 7.03 | 0.00 |
| 39 | 6.46 | 39 | 6.48 | -0.02 |
| 40 | 7.17 | 40 | 7.24 | -0.07 |
| 41 | 7.16 | 41 | 7.19 | -0.03 |
| 42 | 5.62 | 42 | 5.66 | -0.04 |
| 43 | | 43 | | 0.00 |
| 44 | 0.91 | 44 | 0.93 | -0.02 |
| 45 | 1.67 | 45 | 1.68 | -0.01 |
| 46 | 0.87 | 46 | 0.93 | -0.06 |
| 47 | 0.83 | 47 | 0.86 | -0.03 |

**Table S4.** NMR spectroscopic data of neosorangioside A (**3**) acquired at 700/175 MHz in methanol-*d₄*.

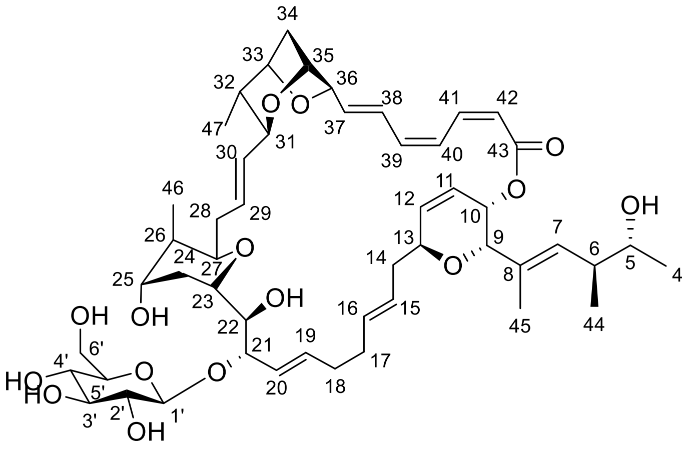


| Position | δC[a] [ppm], type | δH[b] [ppm], mult | COSY[c] | ROESY[d] | HMBC[e] |
|---|---|---|---|---|---|
| 4 | 19.2, CH3 | 1.00, d (6.2) | 5 | 5, 6, 7, 44 | 5, 6 |
| 5 | 71.7, CH | 3.55, qd (6.4, 6.4, 6.4, 4.4) | 4, 6 | 4, 6, 7, 44 | 4, 6, 7, 44 |
| 6 | 40.2, CH | 2.53, dqd (10.0, 6.7, 6.7, 6.7, 4.3) | 44, 5, 7, 44 | 4, 5, 44, 45 | 4, 5, 7, 8, 44 |
| 7 | 130.1, CH | 5.36, m | 6, 45 | 4, 5, 9, 44 | 5, 6, 8, 9, 44, 45 |
| 8 | 133.3, C | | | | |
| 9 | 74.3, CH | 4.23, s | 10, 45 | 7, 10, 14a, 45 | 7, 8, 10, 13, 45 |
| 10 | 66.9, CH | 5.35, dd (5.8, 1.7) | 9, 11 | 9, 11, 45 | 11, 12, 14, 43 |
| 11 | 124.0, CH | 6.03, ddd (9.9, 5.8, 2.2) | 10, 12, 13 | 10 | 10, 12, 13 |
| 12 | 137.2, CH | 6.14, dd (10.1, 3.0) | 11, 13 | 13, 14b | 10, 11, 13 |
| 13 | 75.2, CH | 4.42, dq (11.2, 3.1, 3.1, 3.1) | 11, 12, 14a, 14b | 12, 14b, 15, 16 | 9, 11, 12, 14, 15 |
| 14a | 35.5, CH2 | 2.38, ddd (14.0, 10.9, 8.0) | 13, 14b, 15 | 9, 14a, 15, 16 | 12, 13, 15, 16 |
| 14b | | 2.14, m | 13, 14a, 15 | 12, 13, 14a, 15, 16 | |
| 15 | 128.5, CH | 5.54, ddd (15.3, 8.4, 5.2) | 14a, 14b, 16 | 13, 14a, 14b, 17 | 13, 14, 16, 17 |
| 16 | 134.1, CH | 5.51, ddd (15.3, 7.3, 4.3) | 15, 17a, 17b | 13, 14a, 14b | 14, 17 |
| 17a | 34.1, CH2 | 2.15, m | 16, 17b, 18b | 17b, 20 | 15, 16, 18, 19, 20 |
| 17b | | 2.06, m | 16, 17a, 18a | 15, 17a, 19 | 15, 18, 19 |
| 18a | 35.1, CH2 | 2.15, m | 17b, 18b, 19, 20 | 18b, 20 | 15, 16, 17, 19, 20 |
| 18b | | 2.05, m | 18a, 19, 20 | 18a, 19 | 16, 17, 19, 20 |
| 19 | 136.4, CH | 5.78, ddd (15.3, 7.7, 5.6) | 18a, 18b, 20 | 17b, 18b, 21 | 17, 18, 21 |
| 20 | 127.1, CH | 5.61, br d (9.5) | 18a, 18b, 19, 21 | 17a, 18a, 21, 23 | 18, 21, 22 |
| 21 | 83.3, CH | 4.34, dd (8.4, 3.0) | 20, 22 | 19, 20, 22, 23 | 19, 20, 1' |
| 22 | 75.6, CH | 3.68, m | 21, 23 | 1', 21, 24 | 20, 21, 23, 24 |
| 23 | 74.3, CH | 3.60, ddd (11.5, 8.5, 2.2) | 22, 24a, 24b | 20, 21, 24a, 27 | 21, 22, 25 |
| 24a | 31.6, CH2 | 1.79, br d (14.0) | 23, 24b, 25 | 23, 24b, 25 | 25, 26 |
| 24b | | 1.60, ddd (14.3, 11.7, 2.9) | 23, 24a, 25 | 22, 24a, 25, 46 | 23 |
| 25 | 71.2, CH | 3.85, m | 24a, 24b, 26 | 24a, 24b, 26, 46, 47 | 23, 46 |
| 26 | 38.0, CH | 1.54, m | 25, 27, 46 | 25, 27, 30, 46, 47 | 24, 25, 46 |
| 27 | 74.6, CH | 3.81, m | 26, 28a, 28b | 23, 26, 28a, 30 | 28, 46 |
| 28a | 37.2, CH2 | 2.26, ddd (13.6, 5.0, 2.2) | 27, 28b, 30 | 27, 28b, 29 | 26, 27, 29 |
| 28b | | 2.18, m | 27, 28a, 29, 30 | 28a, 29, 46 | 27, 29 |
| 29 | 132.6, CH | 5.38, m | 28b, 31 | 28a, 28b, 31, 32, 47 | 28, 31 |
| 30 | 133.1, CH | 5.38, m | 28a, 28b, 31 | 26, 27, 31, 46 | |
| 31 | 81.3, CH | 3.85, m | 29, 30, 32 | 29, 30, 37, 46, 47 | 29, 32. 47 |
| 32 | 42.2, CH | 1.42, dq (9.6, 6.8, 6.8, 6.8) | 31, 33, 47 | 29, 33, 34b, 47 | 30, 31, 33, 34, 47 |

| | | | | | |
|---|---|---|---|---|---|
| 33 | 81.3, CH | 4.31, d (6.7) | 32, 34a, 34b | 32, 34a, 34b, 47 | 31, 34, 35, 36, 47 |
| 34a | 39.8, CH2 | 2.04, m | 33, 34b, 35 | 33, 34b, 35, 36 | 31, 32, 33 |
| 34b | | 1.93, dd (11.5, 1.4) | 33, 34a, 35 | 32, 33, 34a, 35 | 32, 35, 36 |
| 35 | 77.7, CH | 4.41, br m | 34a, 34b, 36 | 34a, 34b, 36, 37 | 31 |
| 36 | 82.1, CH | 4.60, br q (2.6, 2.6, 2.6) | 35, 37, 38 | 34a, 35, 37, 38 | 37, 38 |
| 37 | 135.7, CH | 6.25, dd (15.3, 3.7) | 36, 38, 39 | 31, 35, 36, 39 | 35, 36, 38, 39 |
| 38 | 127.5, CH | 7.04, ddd (15.1, 11.5, 1.6) | 36, 37, 39 | 36, 41 | 36, 39, 40 |
| 39 | 138.1, CH | 6.46, m | 37, 38, 40, 42 | 37, 40 | 37, 38, 41, 43 |
| 40 | 126.6, CH | 7.16, m | 39 | 39 | 38, 42, 43 |
| 41 | 139.2, CH | 7.16, m | 42 | 38, 42 | 39, 43 |
| 42 | 119.7, CH | 5.61, m | 39, 41 | 41 | 40, 43 |
| 43 | 167.8, C | | | | |
| 44 | 15.6, CH3 | 0.90, d (6.9) | 6 | 4, 5, 6, 7, 45 | 5, 6, 7 |
| 45 | 14.4, CH3 | 1.66, d (0.9) | 7, 9 | 6, 9, 10, 44 | 6, 7, 8, 9, 10, 44 |
| 46 | 10.8, CH3 | 0.87, d (7.1) | 26 | 24b, 25, 26, 28, 30, 31 | 25, 26, 27 |
| 47 | 15.7, CH3 | 0.83, d (6.7) | 32 | 25, 26, 29, 31, 32, 33 | 31, 32 |
| 1' | 102.8, CH | 4.36, d (7.7) | 2' | 22, 3', 5' | 21, 5' |
| 2' | 75.4, CH | 3.21, dd (9.0, 7.7) | 1', 3' | 3' | 1', 3', 4' |
| 3' | 78.2, CH | 3.35, t (9.0, 9.0) | 2', 4' | 1', 2', 5' | 1', 2', 4' |
| 4' | 71.6, CH | 3.32, d (9.3) | 3', 5' | | 3', 6' |
| 5' | 78.0, CH | 3.22, m | 4', 6'a, 6'b | 1', 3', 6'a, 6'b | 1', 4', 6' |
| 6'a | 62.8, CH2 | 3.82, m | 5', 6'b | 5', 6'b | 4', 5' |
| 6'b | | 3.68, m | 5', 6'a | 5', 6'a | 4', 5' |

[a] Acquired in methanol-$d_4$ at 176.1 MHz and calibrated to solvent signal at 49.2 ppm.
[b] Acquired in methanol-$d_4$ at 700.4 MHz and calibrated to solvent signal at 3.31 ppm.
[c] Proton showing COSY correlations to indicated proton.
[d] Proton showing ROESY correlations to indicated proton.
[e] Proton showing HMBC correlations to indicated carbon.

**Table S5.** $^{13}C$ NMR comparison neosorangioside A (**2**) and sorangioside A (**4**) in methanol-$d_4$.

| Position | δ $^{13}C$ [ppm] Neosorangiosid A (2) | Position | δ $^{13}C$ [ppm] Sorangiosid A (4) | Δ (2-4) |
|---|---|---|---|---|
| 4 | 19.2 | 4 | 28.2 | -9.0 |
| 5 | 71.7 | 5 | 38.5 | 33.2 |
| 6 | 40.2 | 6 | 32.9 | 7.3 |
| 7 | 130.1 | 7 | 134.3 | -4.2 |
| 8 | 133.3 | 8 | 131.3 | 2.0 |
| 9 | 74.3 | 9 | 74.4 | -0.1 |
| 10 | 66.9 | 10 | 66.9 | 0.0 |
| 11 | 124.0 | 11 | 123.9 | 0.1 |
| 12 | 137.2 | 12 | 136.9 | 0.3 |
| 13 | 75.2 | 13 | 76.3 | -1.1 |
| 14 | 35.5 | 14 | 35.3 | 0.2 |
| 15 | 128.5 | 15 | 129.0 | -0.5 |
| 16 | 134.1 | 16 | 132.9 | 1.2 |
| 17 | 34.1 | 17 | 33.6 | 0.5 |
| 18 | 35.1 | 18 | 33.4 | 1.7 |
| 19 | 136.4 | 19 | 136.4 | 0.0 |
| 20 | 127.1 | 20 | 127.2 | -0.1 |
| 21 | 83.3 | 21 | 83.2 | 0.0 |
| 22 | 75.6 | 22 | 76.0 | -0.4 |
| 23 | 74.3 | 23 | 74.0 | 0.3 |
| 24 | 31.6 | 24 | 31.4 | 0.2 |
| 25 | 71.2 | 25 | 71.0 | 0.2 |
| 26 | 38.0 | 26 | 38.0 | 0.0 |
| 27 | 74.6 | 27 | 74.4 | 0.2 |
| 28 | 37.2 | 28 | 37.1 | 0.0 |
| 29 | 132.6 | 29 | 132.4 | 0.2 |
| 30 | 133.1 | 30 | 133.1 | 0.0 |
| 31 | 81.3 | 31 | 81.2 | 0.1 |
| 32 | 42.2 | 32 | 42.1 | 0.1 |
| 33 | 81.3 | 33 | 81.1 | 0.2 |
| 34 | 39.8 | 34 | 39.8 | 0.0 |
| 35 | 77.7 | 35 | 77.5 | 0.2 |
| 36 | 82.1 | 36 | 82.0 | 0.1 |
| 37 | 135.7 | 37 | 134.9 | 0.8 |
| 38 | 127.5 | 38 | 127.8 | -0.4 |
| 39 | 138.1 | 39 | 137.6 | 0.6 |
| 40 | 126.6 | 40 | 127.0 | -0.4 |
| 41 | 139.2 | 41 | 139.1 | 0.1 |
| 42 | 119.7 | 42 | 119.7 | 0.0 |
| 43 | 167.8 | 43 | 167.7 | 0.2 |
| 44 | 15.6 | 44 | 21.7 | -6.1 |
| 45 | 14.4 | 45 | 14.3 | 0.1 |
| 46 | 10.8 | 46 | 10.6 | 0.2 |
| 47 | 15.7 | 47 | 15.4 | 0.3 |
| 1' | 102.8 | 1' | 102.8 | 0.0 |
| 2' | 75.4 | 2' | 75.2 | 0.2 |
| 3' | 78.2 | 3' | 78.1 | 0.1 |
| 4' | 71.6 | 4' | 71.6 | 0.0 |
| 5' | 78.0 | 5' | 77.8 | 0.2 |
| 6' | 62.8 | 6' | 62.8 | 0.0 |

**Table S6.** Blastp results of the CDS regions in the *nsr* BGC.

| CDS Name | Length [AA] | Closest homologue [Organism of origin] | Identity [%] and query coverage [%] | Accession Nr. |
|---|---|---|---|---|
| **NsrJ** | 469 | SorJ [Sorangium cellulosum] | 90.3; 100 | ADN68485.1 |
| **NsrK** | 408 | SorK [Sorangium cellulosum] | 92.7; 100 | ADN68486.1 |
| **NsrL** | 788 | SorL [Sorangium cellulosum] | 91.4; 88 | ADN68487.1 |
| **NsrM** | 1105 | SorM [Sorangium cellulosum] | 88.8; 100 | ADN68497.1 |
| **NsrN** | 469 | SorN [Sorangium cellulosum] | 94.7; 100 | ADN68488.1 |
| **NsrO** | 617 | SorO [Sorangium cellulosum] | 90.1; 100 | ADN68489.1 |
| **Nsr1** | 59 | hypothetical protein [Micromonosporaceae bacterium] | 61.3; 53 | HKE65492.1 |
| **Nsr2** | 88 | acyl carrier protein [Steroidobacteraceae bacterium] | 62.7; 94 | MGH8260247.1 |
| **NsrQ** | 661 | SorQ [Sorangium cellulosum] | 93.3; 100 | ADN68491.1 |
| **NsrA** | 8254 | too big to be analyzed | | |
| **NsrB** | 4970 | SorB [Sorangium cellulosum] | 91.9; 100 | ADN68477.1 |
| **NsrC** | 2076 | SorC [Sorangium cellulosum] | 90.7; 100 | ADN68478.1 |
| **NsrD** | 3192 | SorD [Sorangium cellulosum] | 92.1; 100 | ADN68479.1 |
| **NsrE** | 5314 | SorE [Sorangium cellulosum] | 91.7; 100 | ADN68480.1 |
| **NsrF** | 424 | SorF [Sorangium cellulosum] | 93.2; 100 | ADN68481.1 |
| **NsrGH** | 6699 | SDR family NAD(P)-dependent oxidoreductase [Pendulispora brunnea] | 69.0; 95 | WP_394849115.1 |
| | | SorH [Sorangium cellulosum] | 92.6; 90 | ADN68483.1 |
| **NsrI** | 2610 | SorI [Sorangium cellulosum] | 91.5; 100 | ADN68484.1 |
| **NsrR** | 383 | SorR [Sorangium cellulosum] | 93.2; 100 | ADN68492.1 |
| **NsrS** | 404 | SorS [Sorangium cellulosum] | 92.3; 100 | ADN68493.1 |
| **NsrT** | 584 | SorT [Sorangium cellulosum] | 86.1; 100 | ADN68494.1 |
| **NsrU** | 461 | MATE family efflux transporter [Sorangium sp.] | 96.5; 100 | HTN89125.1 |
| | | SorU [Sorangium cellulosum] | 83.2; 99 | ADN68498.1 |

**Table S7.** Analysis of domains in the *nsr* BGC in *S. cellulosum* Soce417. Substrate specificities of acyl-transferase (AT) domains are based on antiSMASH[1] predictions, comparison to the AT domains in the *sor* BGC[2] (Figure S3) and fingerprint analysis of acyl hydrolase domains[3]. Functionality of ACP domains was determined based on fingerprint analysis.[4] Analysis of the ketosynthase (KS) domains was done based on phylogeny with the transATor tool[5] and clades with the highest score were listed. The stereochemistry predictions for the dehydratase (DH) domains are based on the orientation of the respective hydroxy-group[4]. The listed stereochemistries for the ketoreductase (KR) domains are based on antiSMASH predictions. Activities of all domains were evaluated based on reported fingerprints of active sites.[4,6] Additional modules compared to the biosynthetic machinery in the *sor* BGC are marked bold.

| Gene | Module | Domain | Characterization/comment |
|---|---|---|---|
| NsrO | Trans-AT | AT_a<br>AT_b | Malonyl-CoA; Hydroxy-malonlyl-CoA, Acyl Hydrolase domain<br>Malonyl-CoA |
| NsrA | 1 | KS<br>DH<br>ACP | Clade 8: unusual starter: AMT/succinate<br>Stereochemistry unclear |
| NsrA | 2 | KS<br>**DH**<br>ACP_a<br>ACP_b | Clade 25: completely reduced<br>Stereochemistry unclear |
| NsrA | 3 | KS<br>DH<br>KR<br>C-MT<br>**ACP_a**<br>ACP_b | Clade 74: $\alpha$-Me reduced/keto/D-OH<br>*E*-configured double bond<br>D-configured hydroxy-group |
| NsrA | 4 | KS<br>KR<br>ACP | Clade 113: double bonds (*E*-configured; (some with $\alpha$-Me)<br>L-configured hydroxy-group |
| NsrA | 5 | KS<br>ACP_a<br>**ACP_b** | Clade 66: $\beta$-L-OH |
| NsrA | 6 | KS<br>KR<br>ACP | Clade 73: exomethylene<br>L-configured hydroxy-group |
| NsrA / NsrB | 7 | KS[a]<br>DH<br>ACP<br>DH[b] | Clade 31: non-elongating (bimodule $\beta$-D-OH)<br><br><br>Inactive |
| NsrB | 8 | KS[a]<br>PS<br>ACP | Clade 76: non-elongating (double bonds) |
| NsrB | 9 | KS<br>DH<br>ACP | Clade 26: pyran/furan rings<br>Stereochemistry unclear |
| NsrB | 10 | KS<br>DH<br>KR<br>ACP | Clade 82: double bonds (mostly *E*-configured)<br>*E*-configured double bond<br>D-configured hydroxy-group |
| NsrB / NsrC | 11 | KS<br>DH<br>**KR**<br>ACP | Clade 25: completely reduced<br>*E*-configured double bond<br>D-configured hydroxy-group |
| NsrC / NsrD | 12 | KS<br>KR<br>C-MT<br>ACP_a | Clade 113: double bonds (*E*-configured; (some with $\alpha$-Me)<br>L-configured hydroxy-group |

| | | | |
|---|---|---|---|
| | | ACP_b | |
| | 13 | KS | Clade 68 α-L-OH/Me-β-D-OH |
| | | KR | L-configured hydroxy-group |
| | | ACP_a | |
| | | **ACP_b** | |
| | 14 | KS | Clade 66: β-L-OH |
| | | KR | L-configured hydroxy-group |
| | | C-MT | |
| | | ACP_a | |
| | | **ACP_b** | |
| | 15 | KS | Clade 74: α-Me reduced/keto/D-OH |
| | | DH | *E*-configured double bond |
| | | PS | |
| | | KR | D-configured hydroxy-group |
| NsrE | | ACP | |
| | 16 | KS | Clade 26: pyran/furan rings |
| | | DH | *E*-configured double bond |
| | | KR | D-configured hydroxy-group |
| | | ACP | |
| | 17 | KS | Clade 82: double bonds (mostly *E*-configured) |
| | | DH | *Z*-configured double bond |
| | | KR | L-configured hydroxy-group |
| | | C-MT | |
| | | ACP | |
| | 18 | KS | Clade_68 α-L-OH/Me-β-D-OH |
| | | **O-MT** | |
| | | ACP | |
| | 19 | KS | Clade 103: β-OMe |
| | | DH | *E*-configured double bond |
| | | PS | |
| NsrGH | | KR | D-configured hydroxy-group |
| | | ACP | |
| | 20 | KS | Clade 26: pyran/furan rings |
| | | DH | *E*-configured double bond |
| | | KR | D-configured hydroxy-group |
| | | ACP | |
| | 21 | KS | Clade 82: double bonds (mostly *E*-configured) |
| | | KR | L-configured hydroxy-group |
| | | ACP | |
| | 22 | KS[a] | Clade 31: non-elongating (bimodule β-D-OH) |
| | | DH | *Z*-configured double bond |
| | | ACP | |
| | 23 | KS | Clade 82: double bonds (mostly *E*-configured) |
| NsrI | | DH | *Z*-configured double bond |
| | | KR | L-configured hydroxy-group |
| | | ACP | |
| | | KS[a] | Clade 76: non-elongating (double bonds) |
| | | DH[b] | Inactive |

[a] KS domains have an incomplete CHH catalytic triad and are therefore inactive.
[b] The DH domains are inactive, indicated by the incomplete **H**xxxGxxxxP motif and the missing aspartic acid in an HPALL**D** motif.

### Supplementary Figures

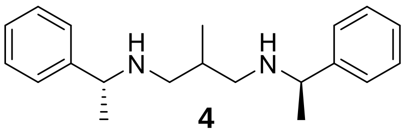


**Figure S1.** Chemical structure of Bis-1,3-methylbenzylamine-2-methylpropane (BMBA-*p*-Me) (**4**).

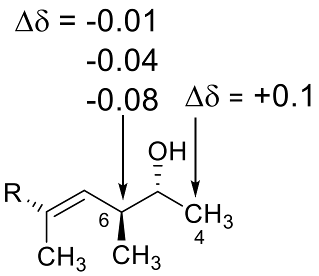


**Figure S2.** $^{13}$C Chemical shift differences of the carbon atoms next to the new stereocenter of neosorangicin A (**1**) in the chiral NMR solvents BMBA-*p*-Me: $\Delta\delta = \delta_{(R,R)\text{-BMBA}} - \delta_{(S,S)\text{-BMBA}}$ in ppm.

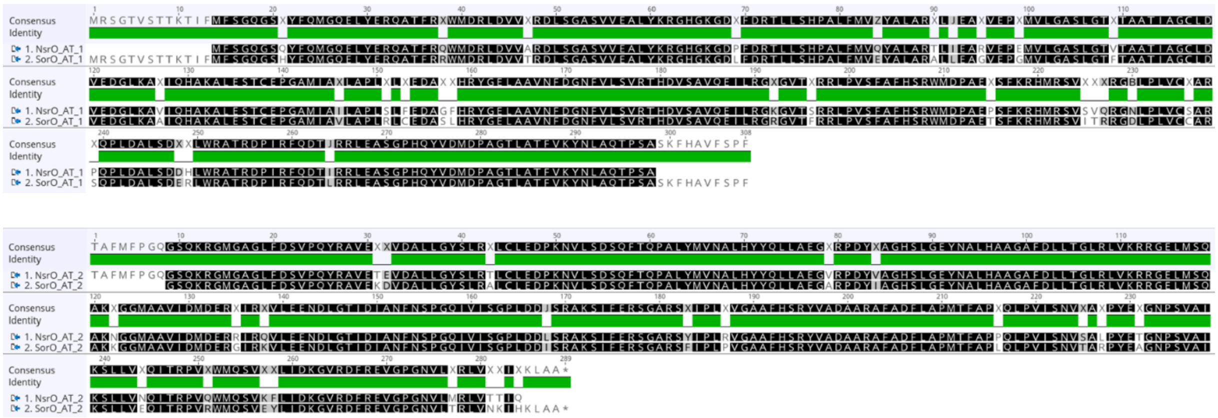

**Figure S3.** Alignment of the AT domains in NsrO in the *nsr* BGC with the ones in SroO in the *sor* BGC. The respective AT domains display a high similarity with 90.2% identity for AT 1 and 91.7% identity for AT 2.

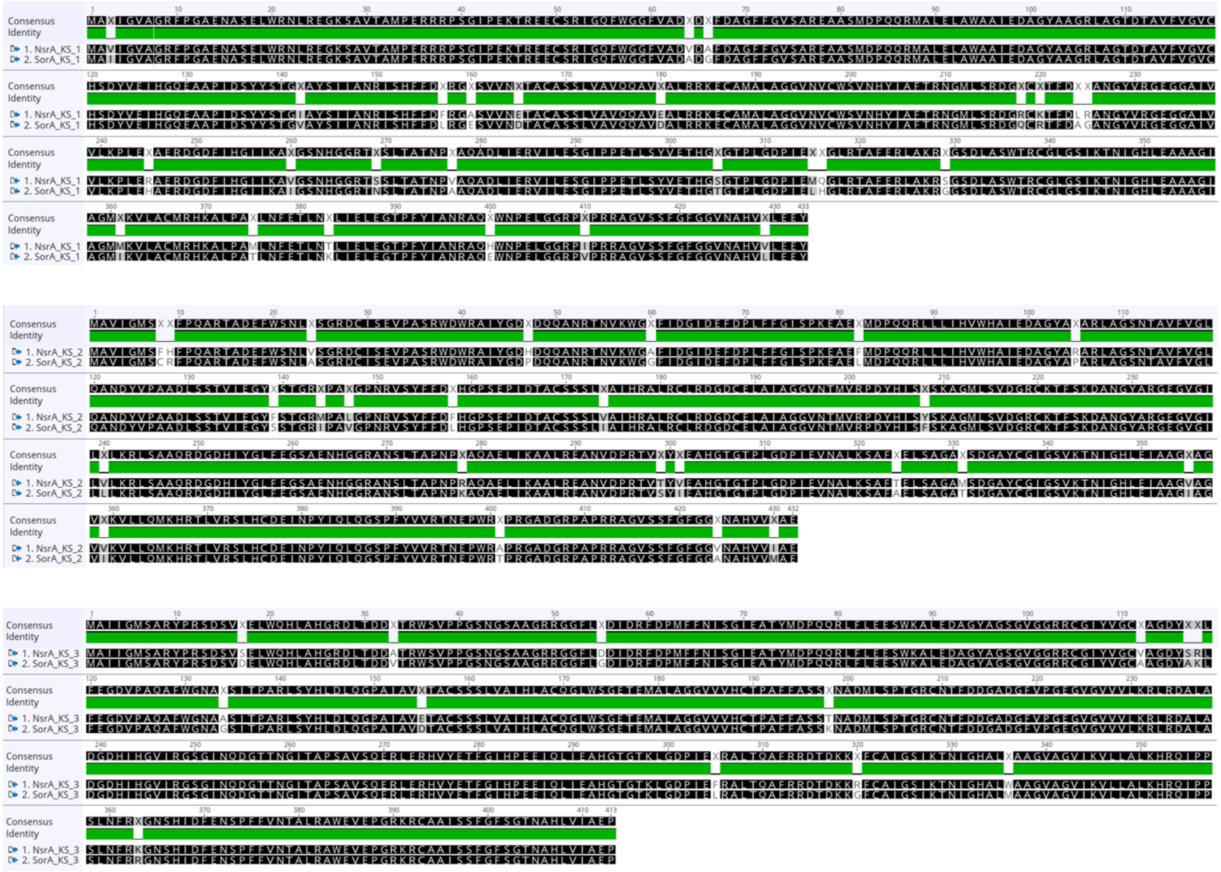

**Figure S4.** Alignment of the KS domains in modules 1-3 in NsrA and SroA. The respective KS domains show a high similarity with 94.0% identity for KS 1, 94.4% for KS 2 and 96.9% for KS 3.

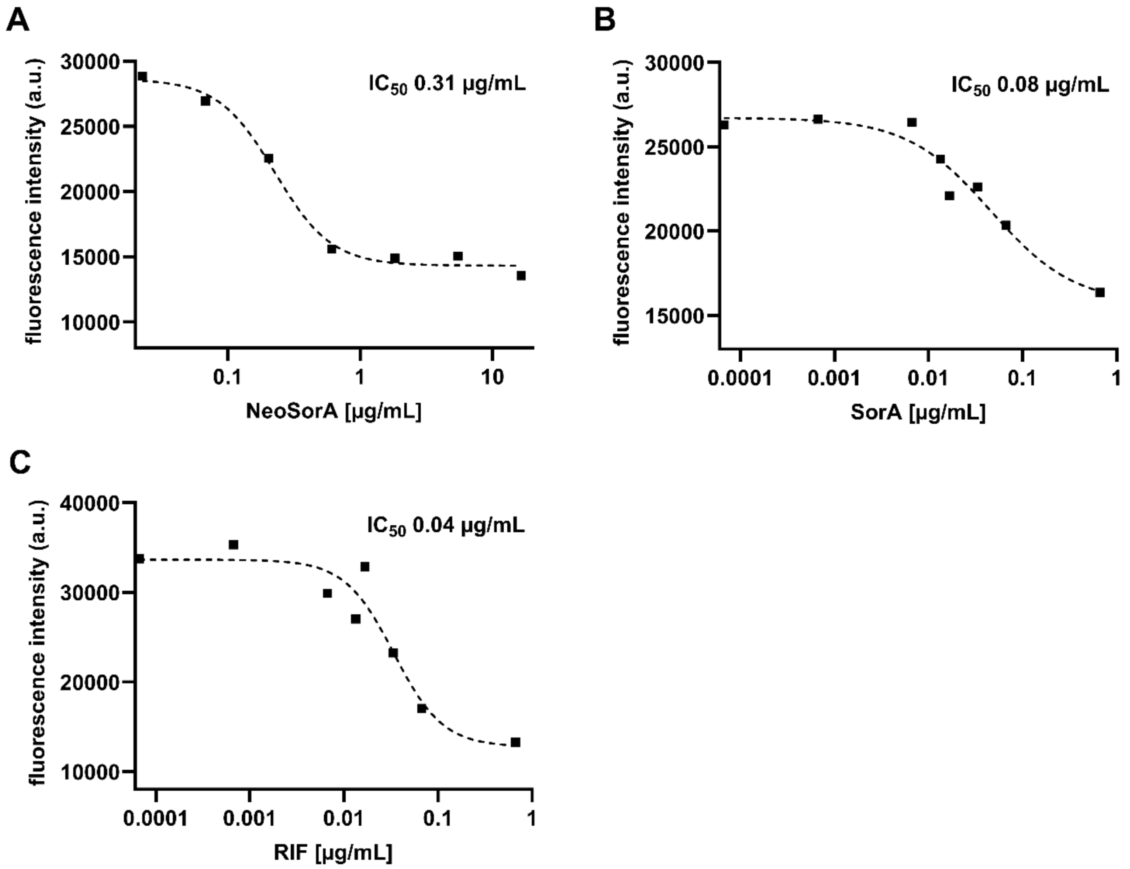


**Figure S5.** *S. aureus* RNA polymerase inhibition assays. Data were plotted using GraphPad Prism (version 10.2.3) and $IC_{50}$ values were determined by sigmoidal curve fitting (**A** neosorangicin A, **B** sorangicin A, **C** rifampicin). NeoSorA: neosorangicin A; RIF: rifampicin; SorA: sorangicin A.

**A**

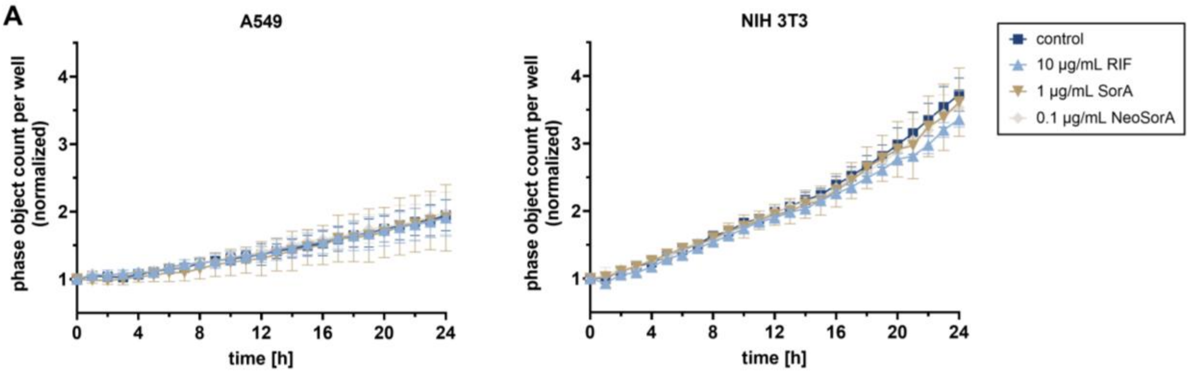


**B**

| | IC$_{50}$ [µg/mL] | | |
|---|---|---|---|
| | 0.1% MeOH | SorA | NeoSorA |
| **A549** | - | 22.5 | 15 |
| **NIH 3T3** | - | 28 | 19 |

**Figure S6. A** Growth curves of A549 and NIH 3T3 cells treated with either 0.1% (*v/v*) methanol, 1 µg/mL sorangicin A (SorA), 0.1 µg/mL neosorangicin A (NeoSorA), or 10 µg/mL rifampicin (RIF). Cells were seeded at $3x10^3$ cells per well in 96-well plates and monitored over 24 hours by live-cell imaging. Object counts were quantified using the Adherent Cell-by-Cell Analysis module and normalized to the initial cell number. Data represent quadruplicate wells. **B** IC$_{50}$ values of the indicated compounds for A549 and NIH 3T3 cells as determined by MTT assay.

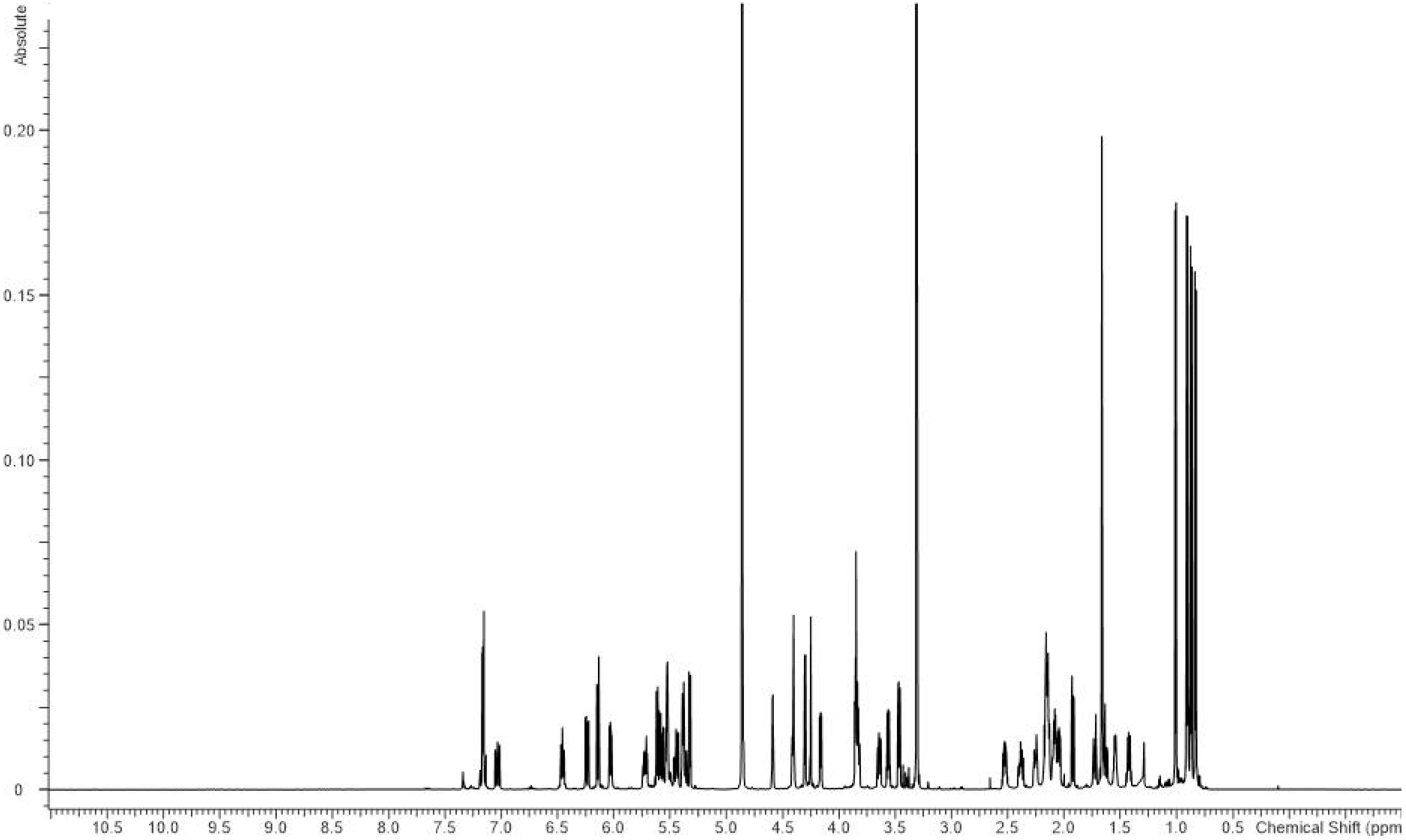


**Figure S7.** $^{1}$H-NMR spectrum of neosorangicin A (**1**) in methanol-$d_4$ (700.4 MHz).

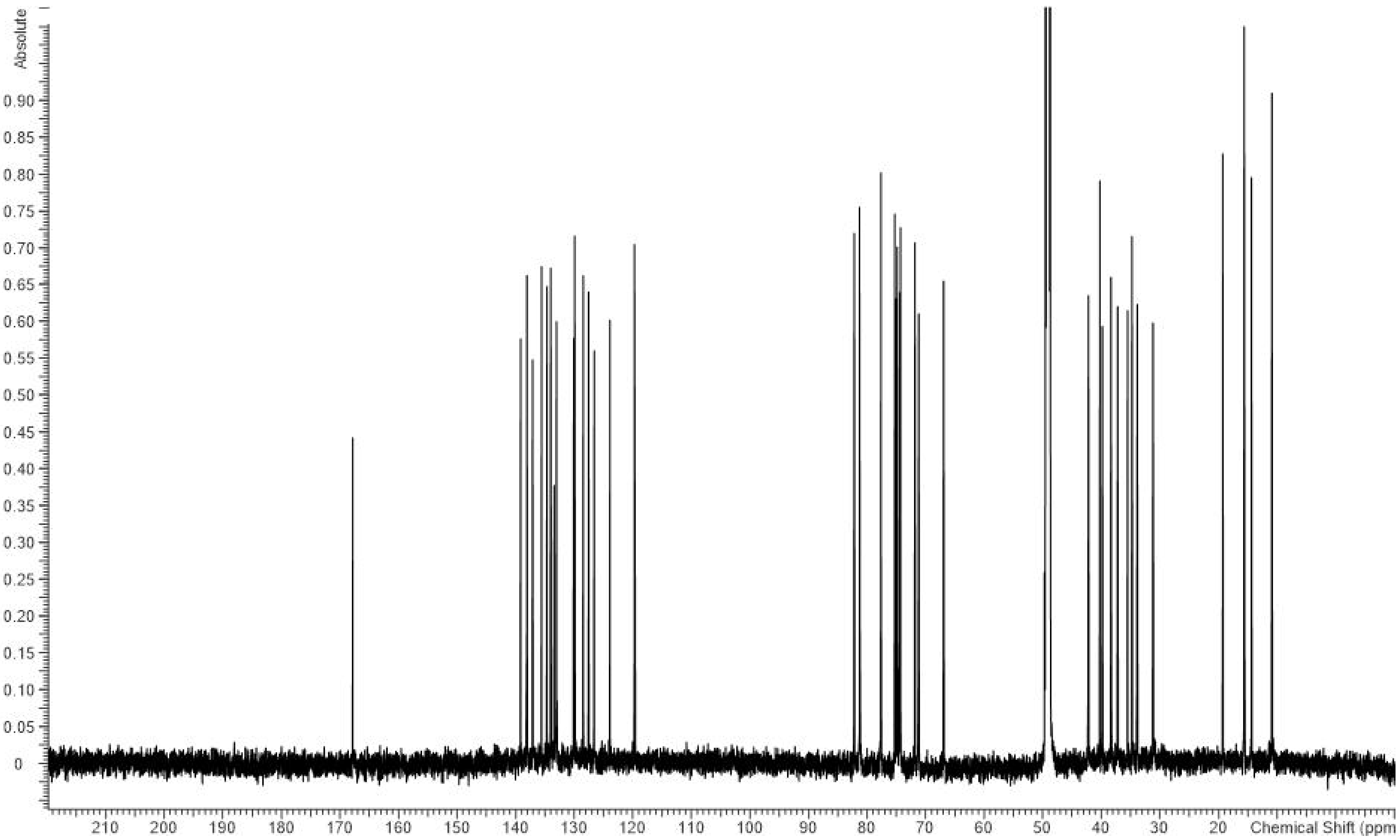


**Figure S8.** $^{13}$C-NMR spectrum of neosorangicin A (**1**) in methanol-*d*$_4$ (176.1 MHz).

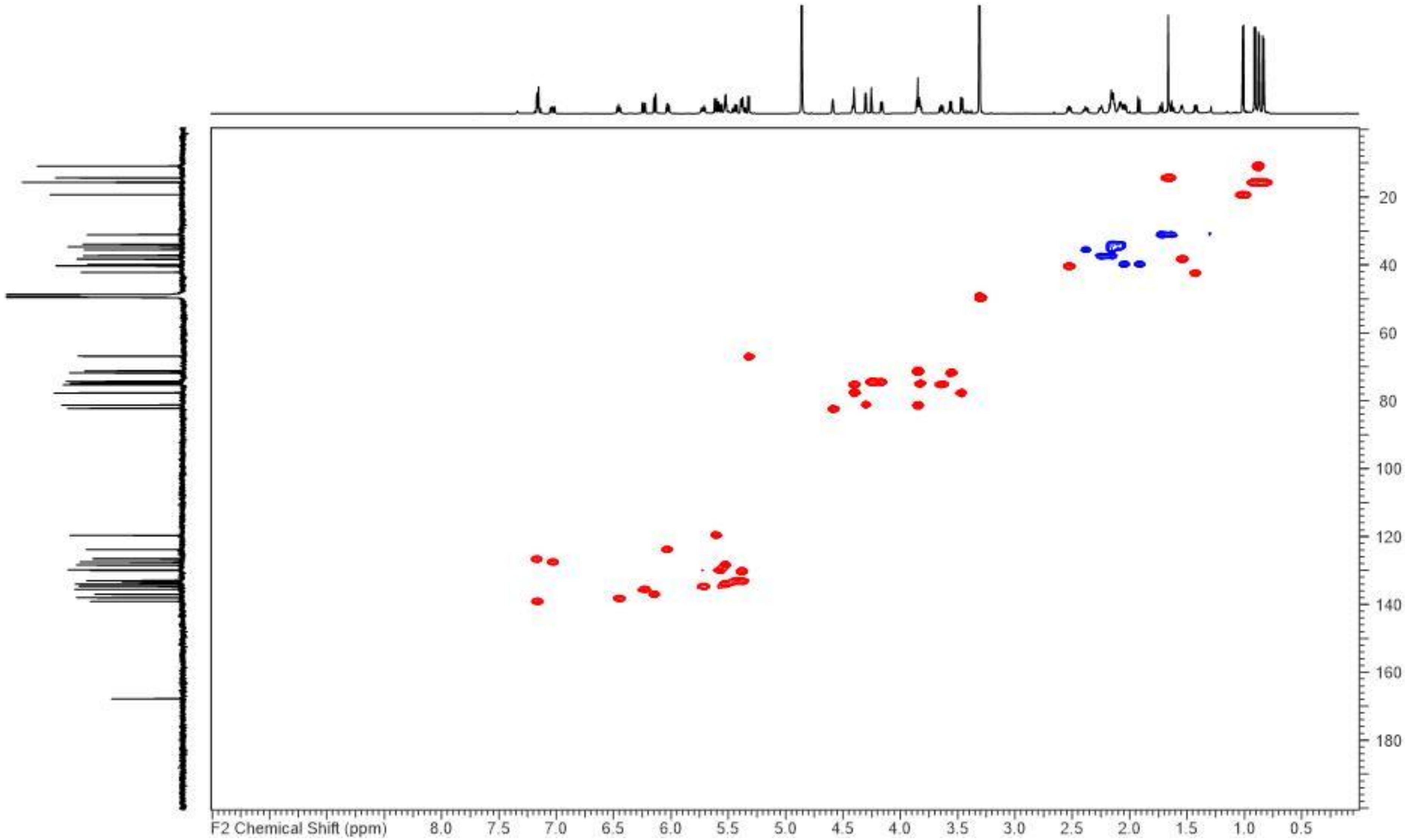


**Figure S9.** HSQC NMR spectrum of neosorangicin A (**1**) in methanol-$d_4$ (176.1/700.4 MHz).

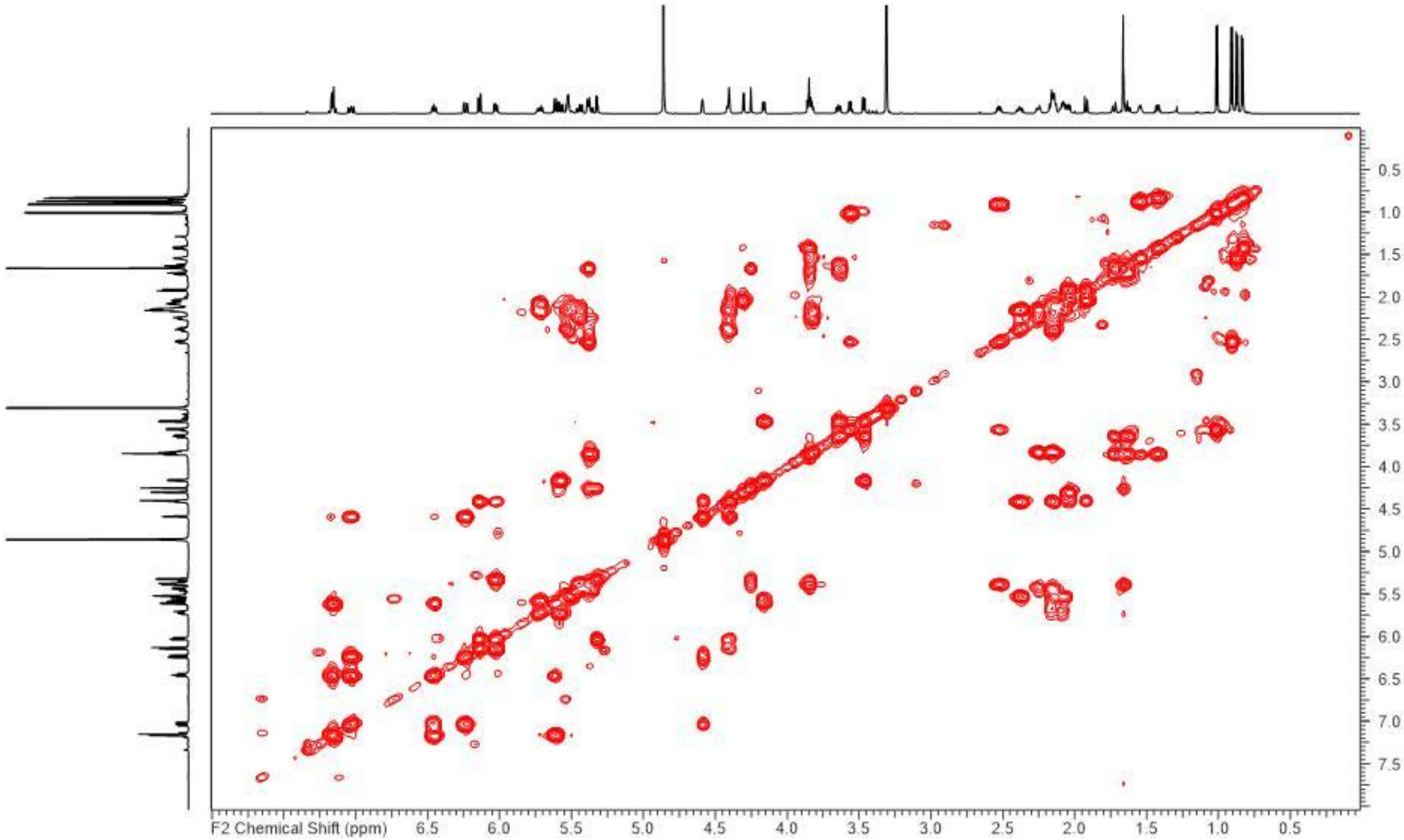


**Figure S10.** COSY NMR spectrum of neosorangicin A (**1**) in methanol-$d_4$ (700.4 MHz).

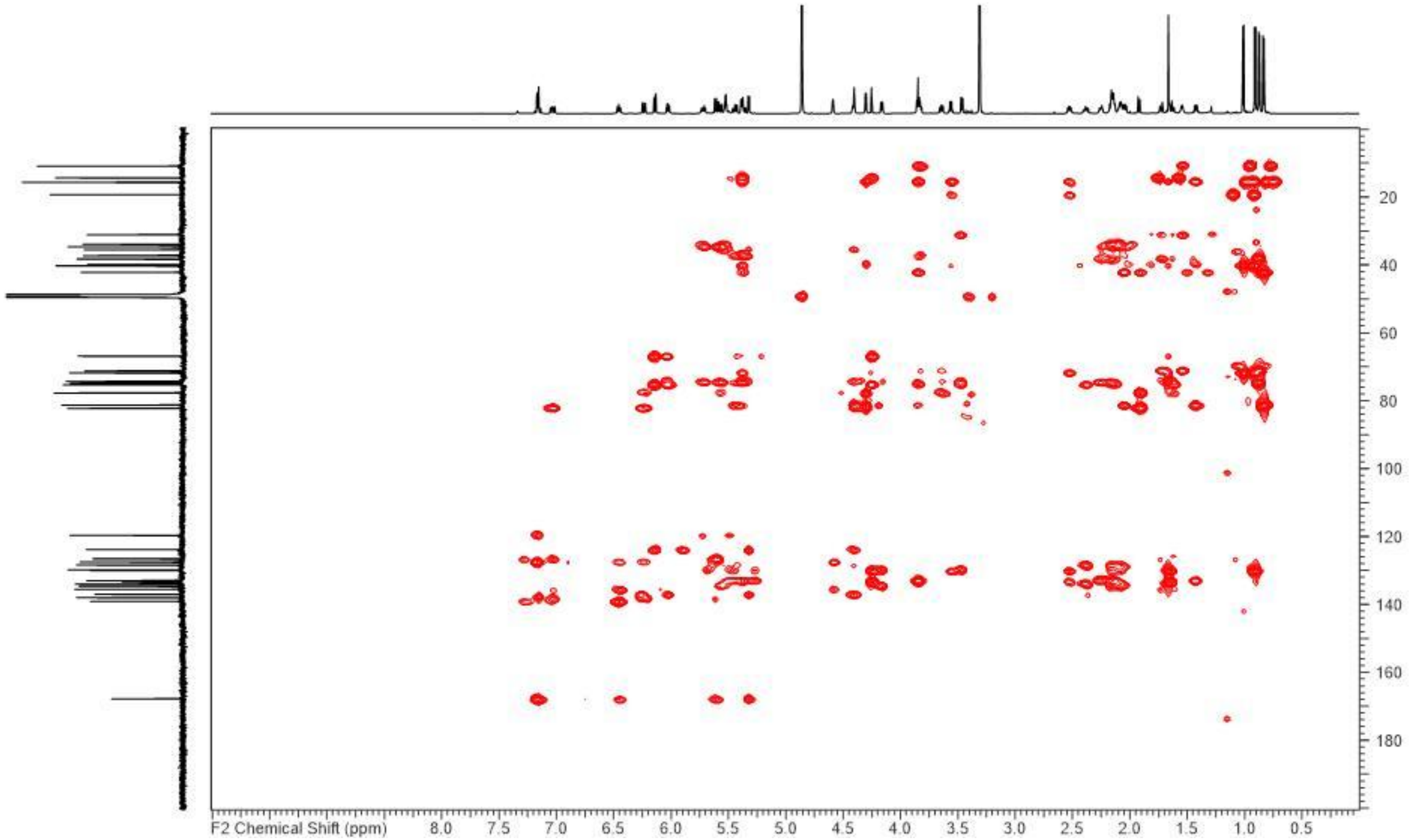


**Figure S11.** HMBC NMR spectrum of neosorangicin A (**1**) in methanol-$d_4$ (176.1/700.4 MHz).

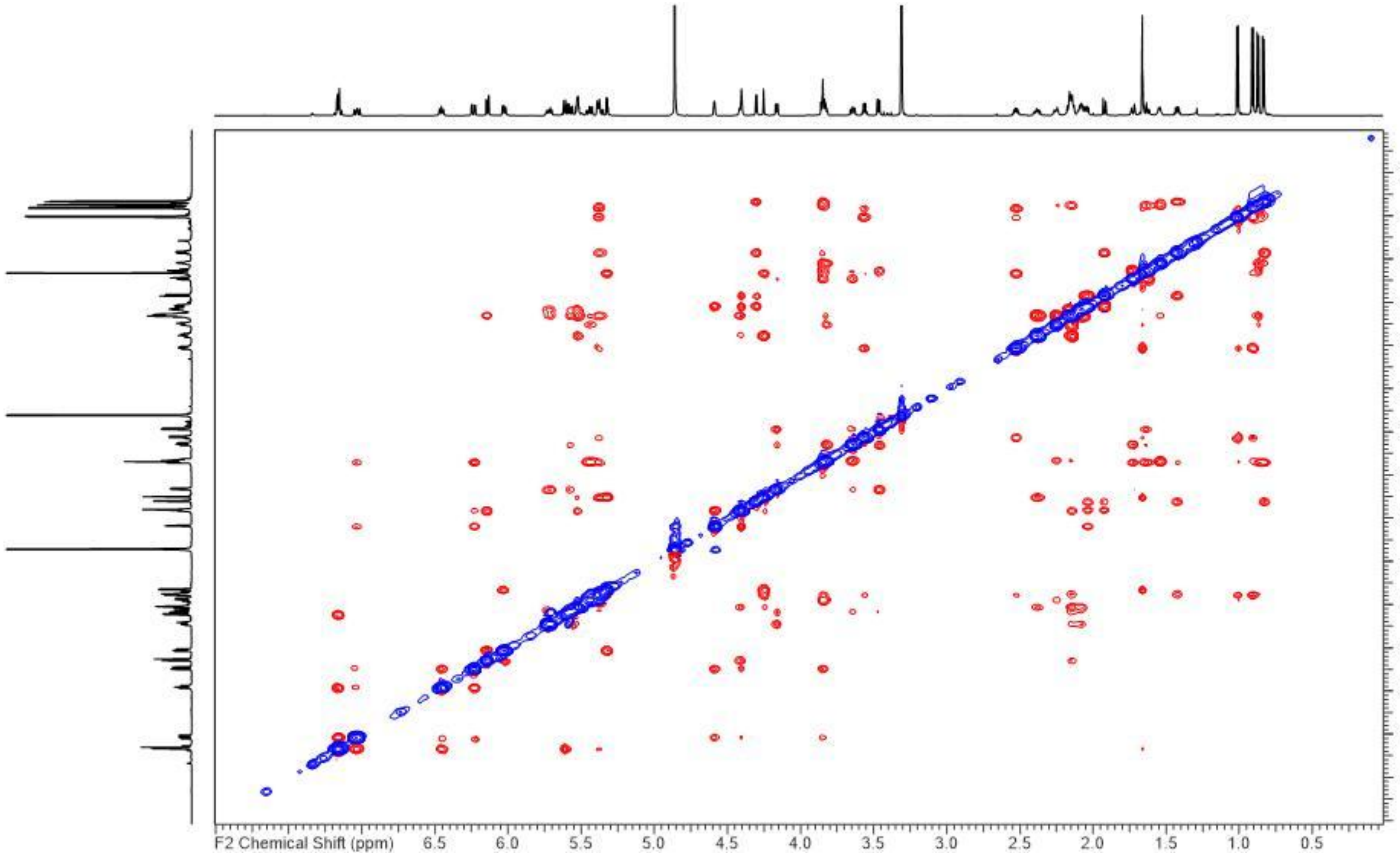


**Figure S12.** ROESY NMR spectrum of neosorangicin A (**1**) in methanol-*d*4 (700.4 MHz).

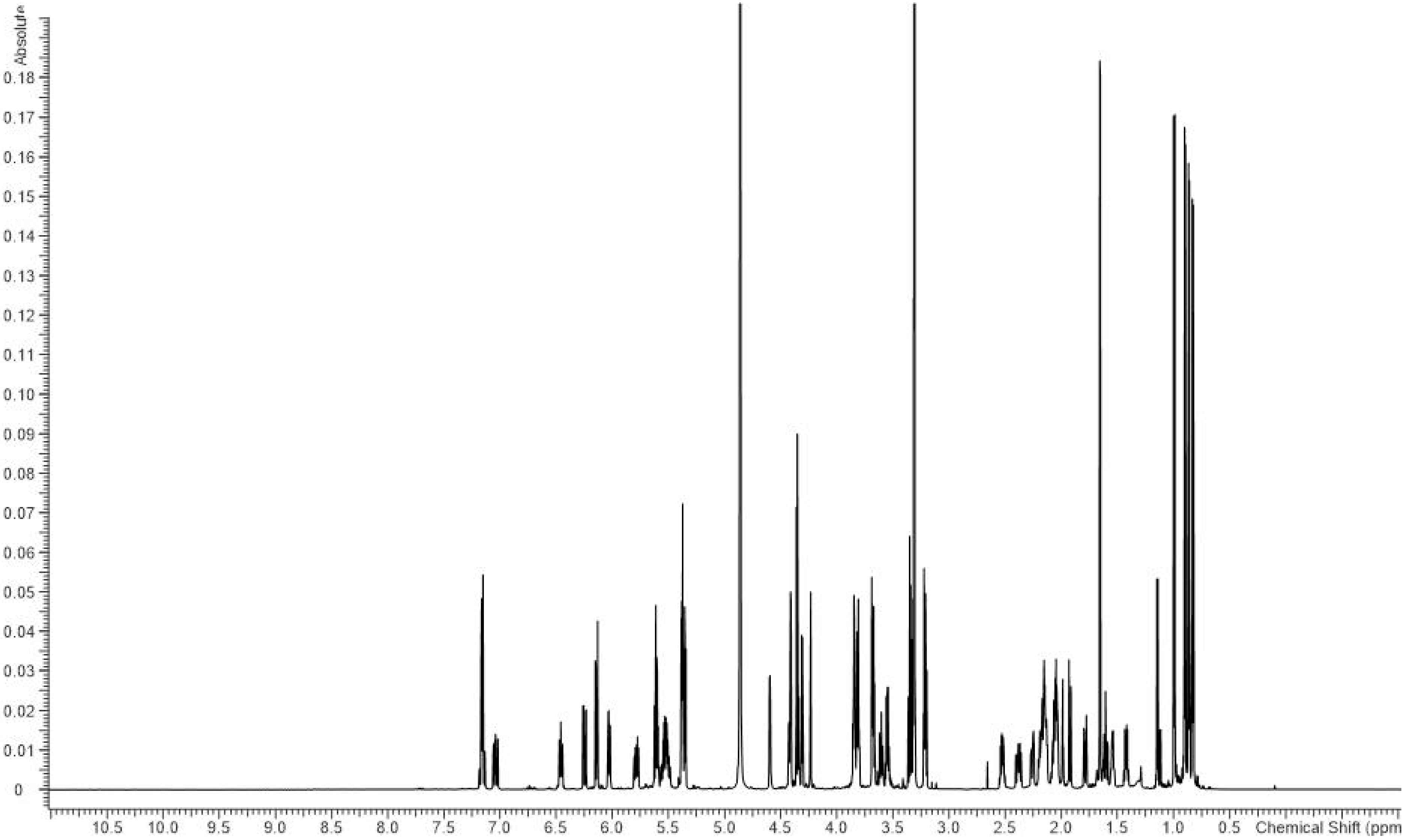


**Figure S13.** $^{1}$H-NMR spectrum of neosorangioside A (**2**) in methanol-$d_4$ (700.4 MHz).

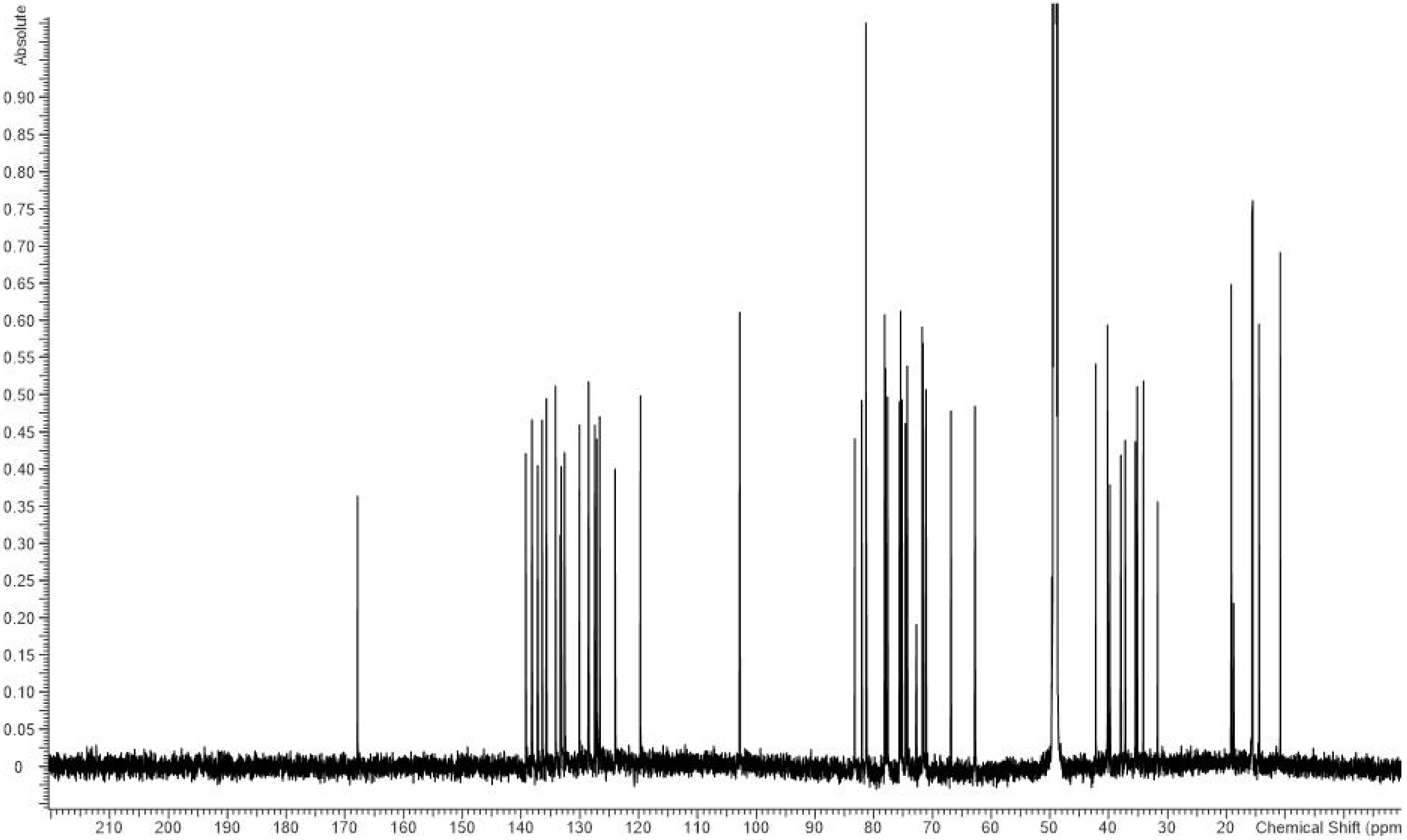


**Figure S14.** $^{13}$C-NMR spectrum of neosorangioside A (**2**) in methanol-$d_4$ (176.1 MHz).

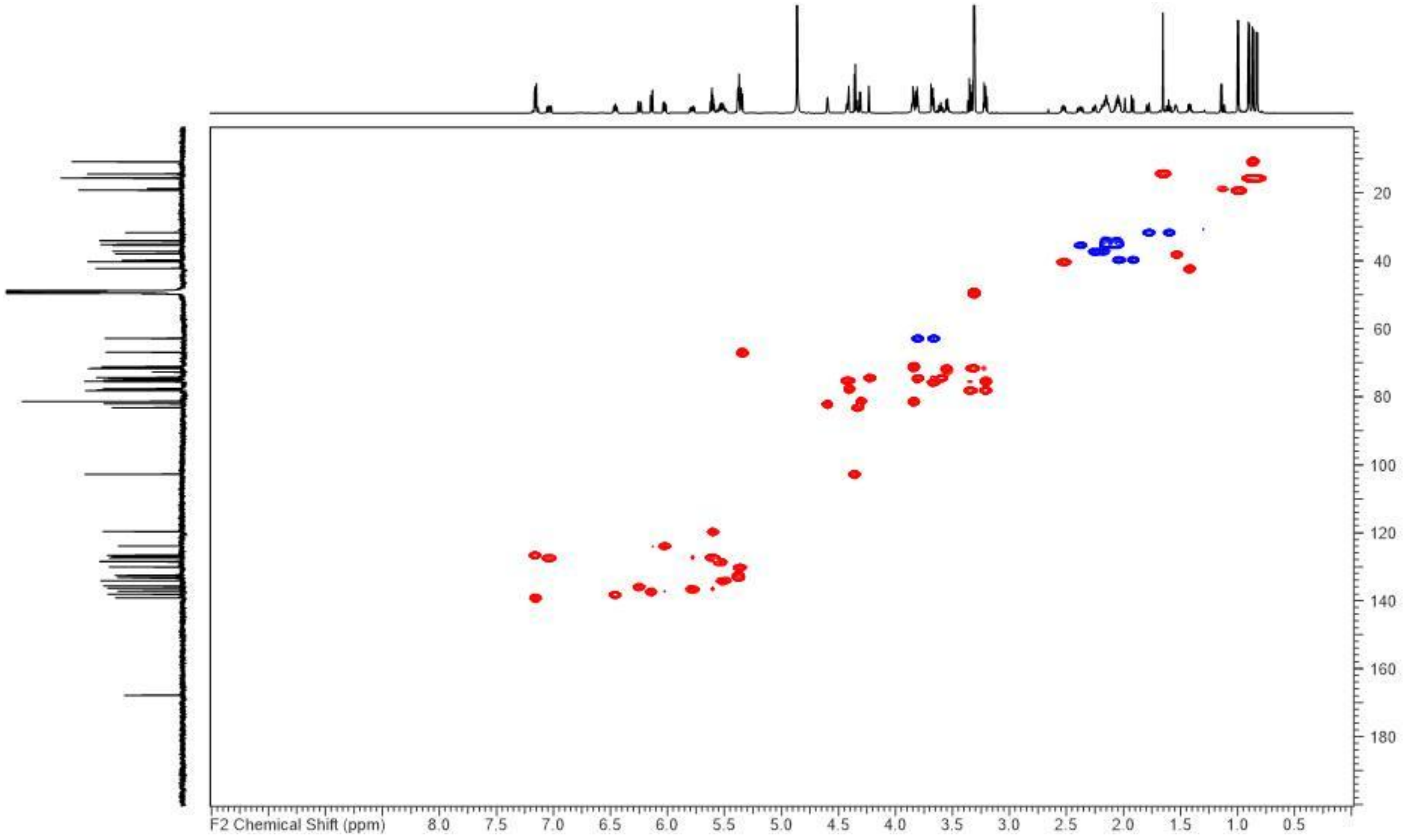


**Figure S15.** HSQC NMR spectrum of neosorangioside A (**1**) in methanol-*d*4 (176.1/700.4 MHz).

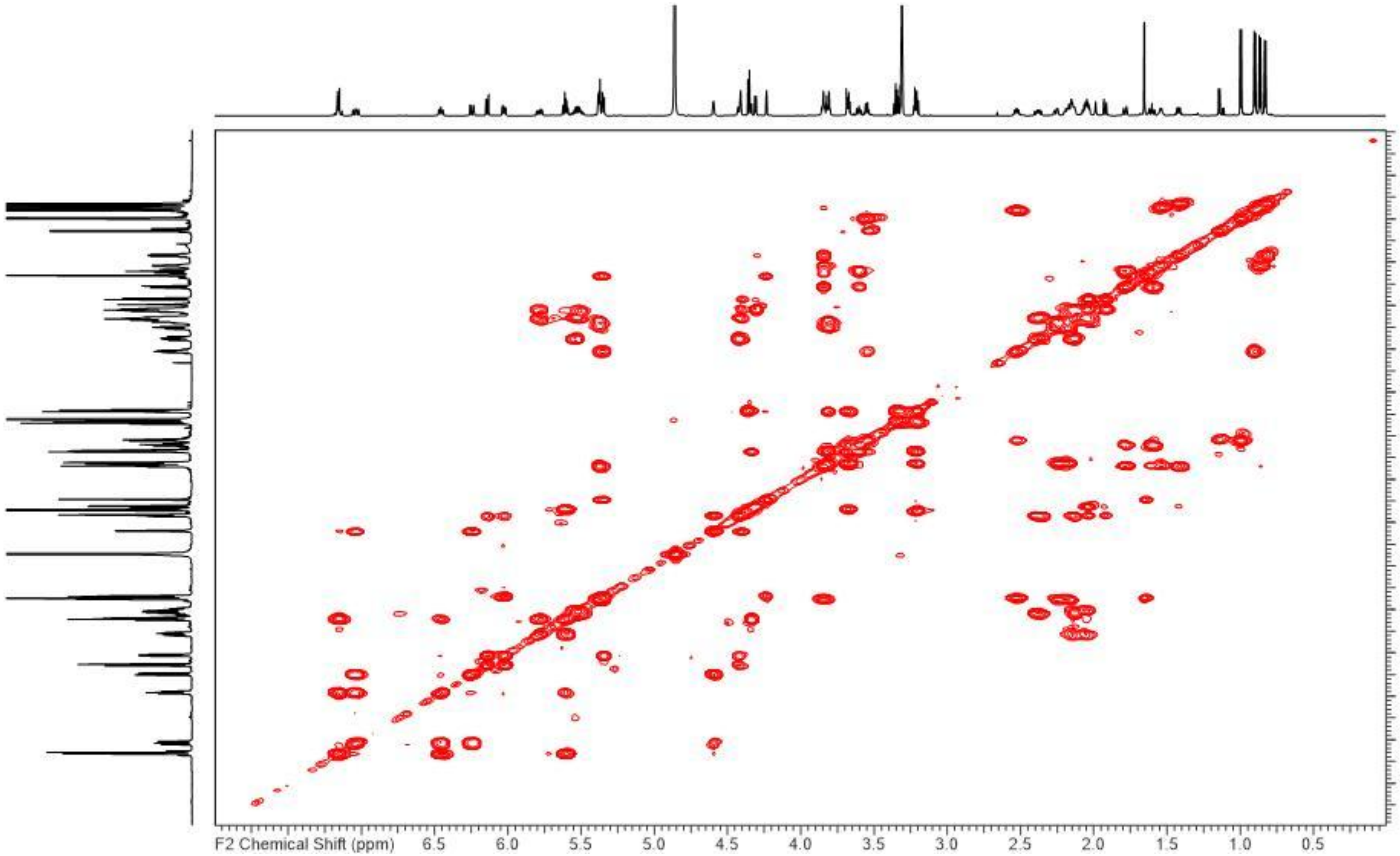


**Figure S16.** COSY NMR spectrum of neosorangioside A (**1**) in methanol-$d_4$ (700.4 MHz).

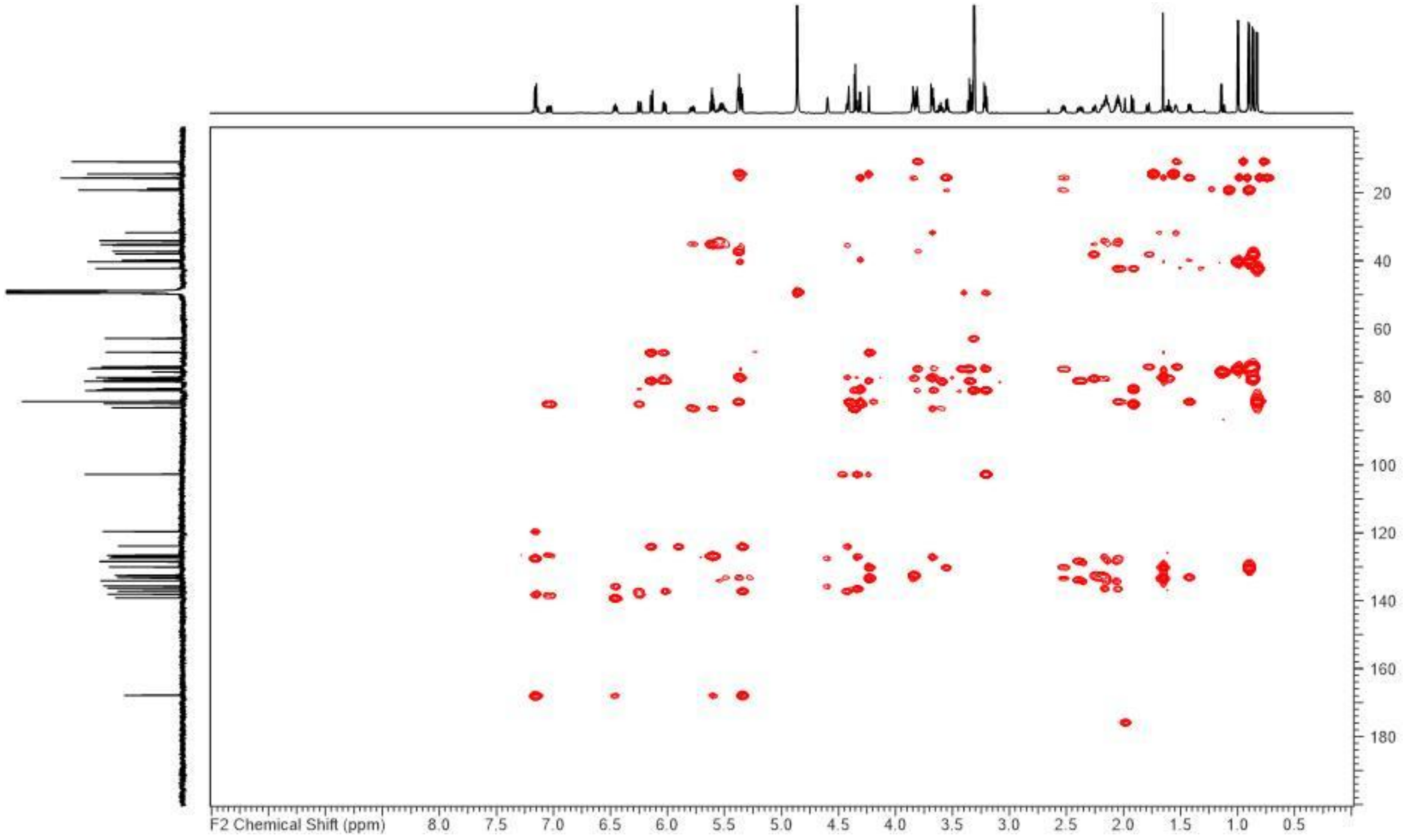


**Figure S17.** HMBC NMR spectrum of neosorangioside A (**1**) in methanol-$d_4$ (176.1/700.4 MHz).

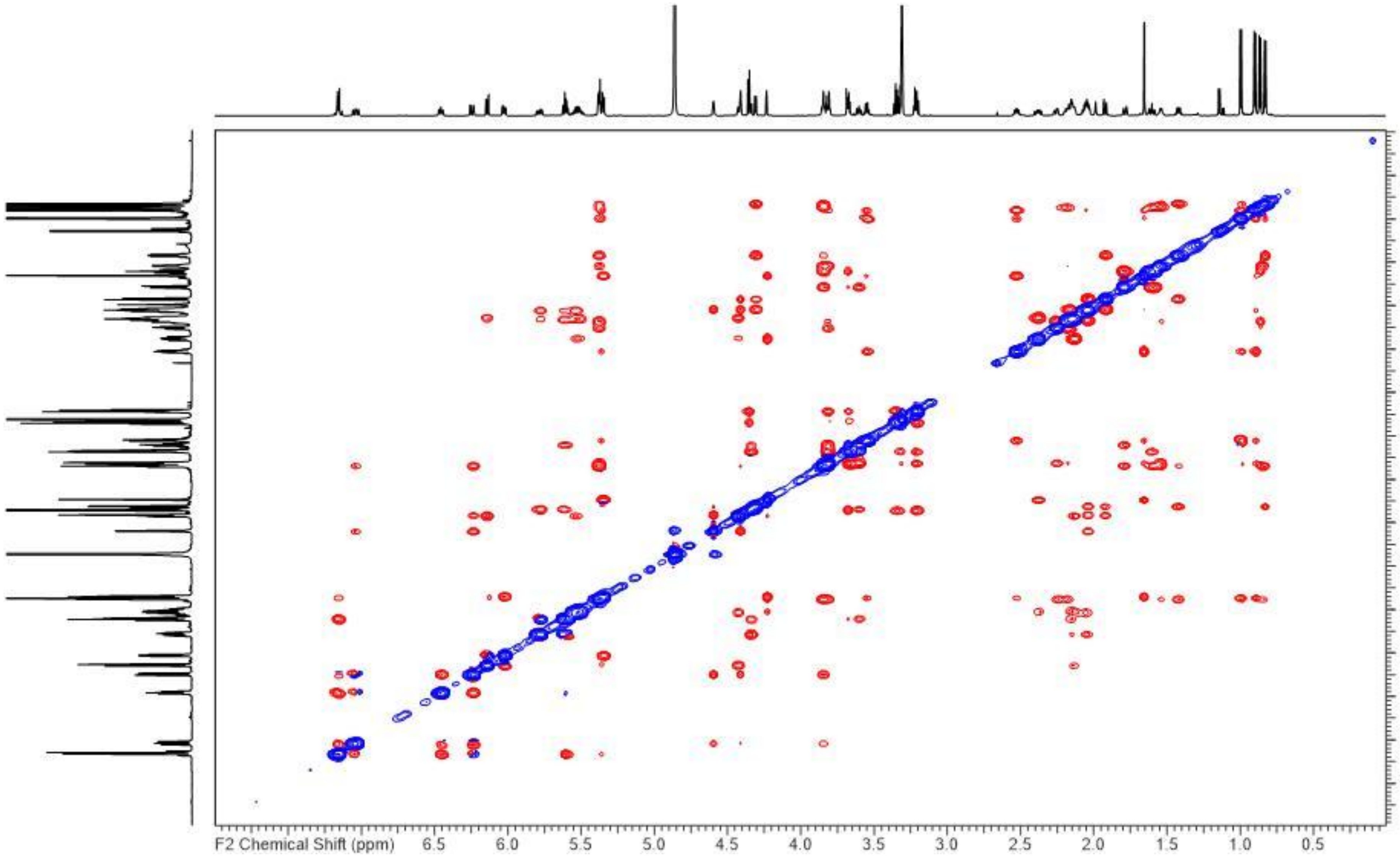


**Figure S18.** ROESY NMR spectrum of neosorangioside A (**1**) in methanol-$d_4$ (700.4 MHz).
